# Impact of the extinct megaherbivore Steller’s sea cow (*Hydrodamalis gigas*) on kelp forest resilience

**DOI:** 10.1101/2022.07.15.500280

**Authors:** Peter D. Roopnarine, Roxanne M. W. Banker, Scott Sampson

## Abstract

Giant kelp forests off the west coast of North America are maintained primarily by sea otter (*Enhydra lutris*) and sunflower sea star (*Pycnopodia helianthoides*) predation of sea urchins. Human hunting of sea otters in historic times, together with a marine heat wave and sea star wasting disease epidemic in the past decade, have devastated these predators, leading to widespread occurrences of urchin barrens. Since the late Neogene, species of the megaherbivorous sirenian *Hydrodamalis* ranged throughout North Pacific giant kelp forests. The last species, *H. gigas*, was driven to extinction by human hunting in the mid-18^th^ century. *H. gigas* was an obligate kelp canopy browser, and its body size implies that it would have had a significant impact on the system. Here we hypothesize that sea cow browsing would have promoted a denser understory algal assemblage than is typical today, thereby providing an alternative food resource for urchins, resulting in enhanced forest resilience. We tested this hypothesis with a mathematical model, comparing historical and modern community responses to marine heat waves, sea star wasting disease, and the presence of sea otters. We found that forest communities were highly resistant to marine heat waves, yet susceptible to sea star wasting disease, and to disease in combination with warming. Resistance was greatest among systems with both sea cows and sea otters present. Most simulations that transitioned to barrens did so temporarily, recovering after about 10 years. Historical communities, however, exhibited delayed transitions after perturbation relative to modern communities and faster recovery times. Sea cow browsing facilitated denser algal understories, enhancing resilience against modern perturbations. We propose that operationalizing these findings by mimicking the ecological impact of sea cow herbivory could enhance kelp forest resilience.

## 1 Introduction

The time frame of concern for most conservation projects is limited to a narrow temporal window, typically spanning less than a century [1]. When contemplating interventions, conservation efforts tend to focus on the recent past and near future. This strategy is problematic in a world where ecosystems have been degenerating for centuries to millennia, and climate is shifting rapidly, resulting in novel ecological conditions. At a minimum, understanding what diversity, integrity, and resilience once looked like for a given ecosystem is critical if we are to be successful in regenerating what might be termed “ecosystem health.” Rigorous exploration of likely future impacts resulting from climate warming and other factors, including sequential reintroduction of species, is essential to increase success probabilities. Here we propose a Past-Present-Future (PPF) approach to conservation rooted in mathematical modeling, illustrating this proposed method with a study of the giant kelp ecosystem in the northeastern Pacific Ocean.

Giant kelp forests are one of the most productive marine ecosystems in the world[2], distributed in cool temperature coastal regions of both hemispheres [3, 4]. Kelp forests of the northern Pacific are particularly iconic, and studies on these ecosystems have become fundamental to the concepts of keystone predators, foundation species, and alternative ecological states [5, 6, 7]. Many kelp-dominated communities, including those engineered by non-gigantic species, include sea urchins as major grazers of the kelp. The kelp-urchin system typically exhibits two states, forest and barrens [7, 8]. Various circumstances can lead to increased or uncontrolled urchin grazing, ultimately resulting in a landscape denuded of kelp. The productivity and biodiversity of such communities are so drastically reduced that the community state is referred to as an “urchin barrens” [9]. The transition from a forest to barrens state is well understood in north Pacific forests: predatory sunflower sea stars (*Pycnopodia helianthoides*) and sea otters (*Enhydra lutris*) control the size of sea urchin populations (purple and red urchins, *Strongylocentrotus purpuratus* and *Mesocentrotus franciscanus*, respectively), facilitating the development of dense kelp forests often dominated by the species *Macrocystis pyrifera* and *Nereocystis luetkeana*. Reduction of either predatory species may be sufficient to drive a transition from the forest to the barrens state, and it has been suggested that predatory suppression of urchins was essential to the diversification and geographic expansion of kelps in the North Pacific during the late Cenozoic [10]. Historical anthropogenic extirpation of the otter in many areas resulted in the conversion of numerous forests to barrens in the 19^th^ and 20^th^ centuries. Recent emergence of sea star wasting disease (SSWD) in the North Pacific, recognized as early as 2013, devastated populations of *P. helianthoides*, releasing the urchins from predatory control and facilitating another widespread transformation of forests to barrens.

### 1.1 The North Pacific kelp-urchin system

The forest state consists of dense concentrations of giant kelp that grow from holdfasts anchored to hard substrate at depths up to 15 m. Robust stalks extend from the substrate to the surface, bearing blade-like fronds that capture light for photosynthesis. In *M. pyrifera*-dominated forests, blades form a dense canopy that block and capture up to 90% of incident sunlight at the surface [11, 12]. Forests with dense canopies have diverse assemblages of other organisms, high overall primary productivity, baffle wave energy and storm surges, and are conventionally considered to be the “healthy” state of kelp forests (Fig. 1A). Adult kelp, however, do inhibit other photosynthesizers, including phytoplankton, other benthic algae and juvenile kelp [13], because of their high light absorption and blocking, as well as their competition for space on the substrate. The barrens, in contrast, consists of a hard substrate dominated numerically by urchins and largely devoid of kelp or understory algae (Fig. 1B). Biodiversity is lower in barrens, probably the result of lowered primary productivity, the loss of a three dimensional foundational structure, and an increase of hydrodynamic energy.

**Figure 1.**
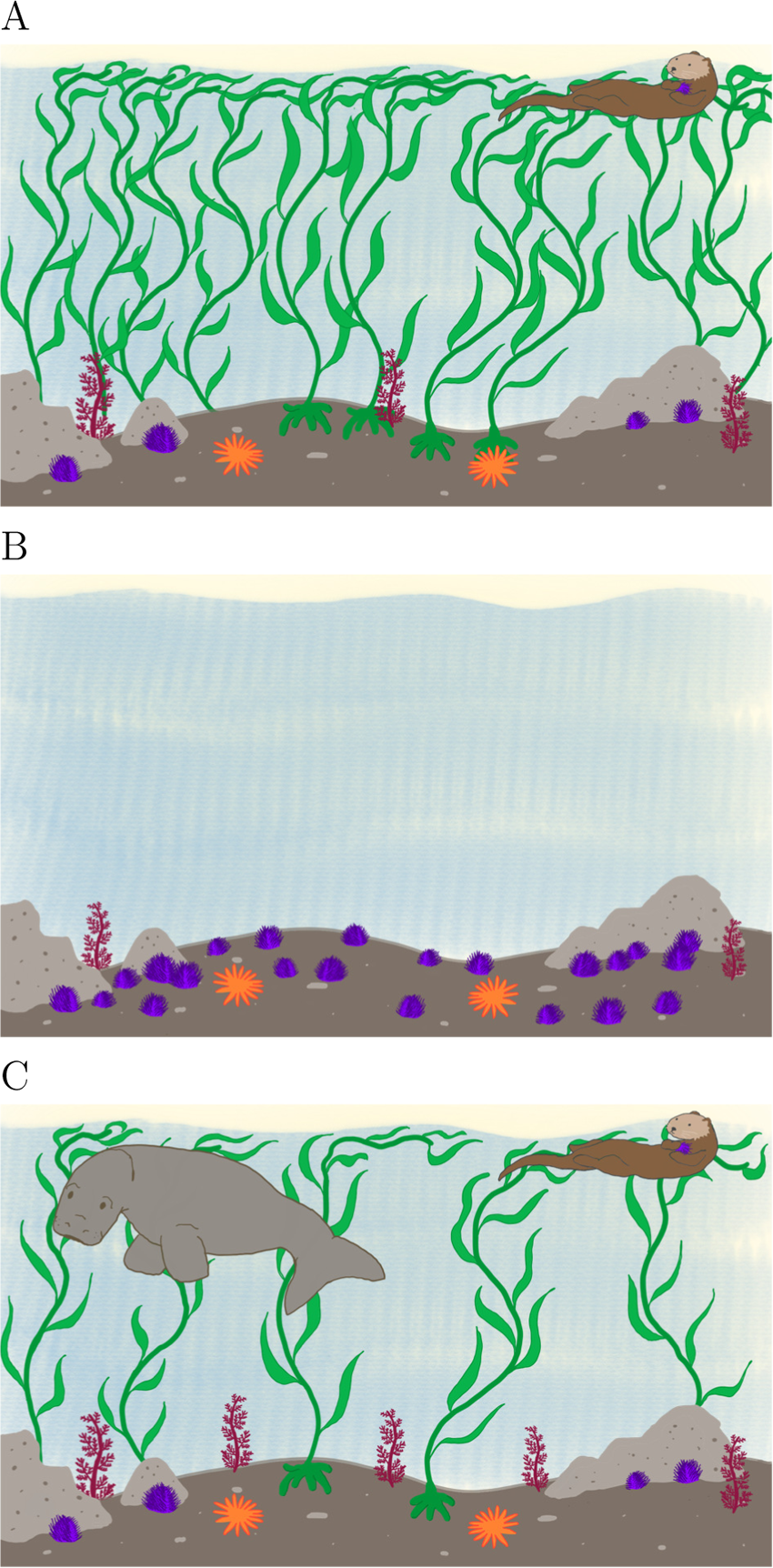

The transition of a forest to the barrens state is often abrupt relative to the duration of the forest, and is preceded by an eruption of urchins that actively forage and prey on adult kelp holdfasts and stipes. Typically, a large fraction of urchins live sheltered and crowded into substrate crevices when predators are present, capturing and consuming the abundant drift detritus produced by the fragmentation of kelp fronds and other macroalgal species. Two factors can increase the fraction of urchins that expose themselves to graze on kelp plants: a reduction of predation pressure or a decline in the availability of drift detritus. Less predation pressure results in growth of the urchin population and a reduction of available crevice space and food availability, while a reduction of drift detritus concentration leads directly to starvation, which may be critical in determining urchin feeding modes [14]. In both cases, urchins will begin to leave crevices in increasing numbers to graze directly on kelp which results in the death of individual kelp. This may create a positive or reinforcing feedback, whereby the loss of adult kelp further reduces the availability of drift detritus, prompting more urchins to become mobile foragers. Furthermore, as kelp density decreases, urchin starvation increases leading to a decline in the nutritive condition of the urchins. Otters will generally avoid predation on less nutritive urchins, further exacerbating and accelerating the transition from forest to the denuded barrens state. Barrens can transition to forest if urchin populations decline sufficiently because of starvation. Furthermore, Feehan et al. [15] have shown that offshore export of particulate kelp detritus is an important food source for urchin larvae. Hence the devastation of kelp productivity by adult urchins could generate a negative feedback on the sustainability of the urchin population. Nevertheless, urchins appear capable of subsisting at low tissue mass for long periods of time under low nutrient conditions, a condition termed “zombie urchins” [16]. Transition to forest can also be facilitated by an increase of sunflower sea star populations, or by exceptionally high and successful recruitment of juvenile kelp [17].

### 1.2 Current state

The dominant state of giant kelp communities in the northern Pacific over the past several decades has been widespread forest, maintained by healthy populations of sunflower sea stars and the return of stable sea otter populations in some areas of their former range. The situation changed dramatically in 2013-2014 with the outbreak of sea star wasting disease (SSWD) on both coasts of North America, and a catastrophic shift to a reduced kelp canopy and dominance of urchin barrens communities [18]. The disease quickly devastated sunflower sea star populations, with the species disappearing entirely in large parts of its range. The rapid decline of the sea stars [19] was followed by widespread “outbreaks” of urchins, with increases in the numbers of actively foraging urchins and apparently local population sizes. These outbreaks resulted in the transition of numerous kelp forests to urchin barrens, e.g. bull kelp (*Nereocystis luetkana*) being reduced by more than 90% [18]. The loss of kelp productivity and alteration of the habitat has had cascading effects throughout forest communities; abalone (*Haliotis*) populations were reduced by 80% by 2017 [18]. That situation that has persisted up to the present, although some recovery of kelp has been noted since summer 2021 along the coast of California [20]. There has been no documented recovery of the sunflower sea stars [19].

An additional factor in the decline of the forests may have been the onshore expansion of a persistent body of warm water, colloquially known as “the blob”, around the same time. This persistently cohesive and anomalously warm body of water, first detected in the sub-polar northern Pacific in late 2013, expanded in 2014 to encompass almost the entire west coast of North America, with sea surface temperatures reaching 2.5-3.0^°^C above average. The marine heat wave (MHW) persisted from 2013-2016 in some coastal areas [21]. It has been speculated that the warm temperatures had a detrimental affect on kelp productivity, thereby triggering urchin outbreaks [18]. Certainly the coincidence of the MHW, SSWD, and the possibility of synergistic effects cannot be overlooked, and the MHW has been suggested as a trigger for SSWD in cooler waters [22].

### 1.3 A paleobiological alternative

The heavily studied and well understood giant kelp ecosystem may be historically young in parts of the North Pacific, at least in its present state. A megaherbivorous mammal, the Steller’s sea cow (*Hydrodamalis gigas*), inhabited kelp ecosystems in the Comander Islands in historical times. This giant sirenian was first described by Georg Wilhelm Steller, a scientist who accompanied Russian commercial voyages to the islands in 1741. The species was quickly exploited as a source of fresh meat and is believed to have become extinct by 1768. Individuals attained very large sizes, up to 9m in length, and are estimated to have weighed up to 10 tons. *H. gigas* was an obligate kelp feeder, apparently incapable of submerging because of its high bouyancy, and therefore browsed the giant kelp canopy. There are no recorded observations of the species beyond the Aleutian Islands, but fossils of both *H. gigas*, and other members of the genus that might represent different species, are known from the Pliocene of California and Baja California, and the post-Pliocene of Japan [23]. The genus may have evolved opportunistically with the late Neogene regional expansion of kelp forests. The rarity (34 partial skeletons have been recorded) of fossil remains makes it impossible to discern whether the genus once encompassed this entire range, including the Arctic during interglacial intervals, or if instead the range was fragmentary, responding perhaps to variable oceanographic conditions after the onset of northern Hemisphere cooling (*∼*2.4 mya). Similarly, previous efforts to split the fossils into a phyletic lineage of multiple species is questionable [23], particularly given a sparsity of specimens, and reliance on overall body size as a primary differentiating trait among the putative species. Nevertheless, all occurrences of *Hydrodamalis*, fossil and historical, imply a dependency on high productivity coastal habitats, and given body size and the angle of the snout, a feeding habit of surface browsing. Several hypotheses have been proposed to explain rarity, geographic restriction, and extinction of *H. gigas*, but none are conclusive (reviewed in Supplementary Material).

The impacts of megaherbivores on community structure and ecosystem functioning has been well established, and their conservation is currently an issue of major concern. Given our understanding that these animals often contribute significantly to the dynamics of their communities, including stability, resilience and state transitions, it is reasonable to speculate that *H. gigas* (and congeneric species) could have played a significant ecological role in kelp forest communities. Recently, Bullen et al. [24] presented six hypotheses of how *H. gigas* may have structured giant kelp communities. Several of the hypotheses predict that grazing of kelp fronds by sea cows opened up the canopy, increasing light intensity in both the water column and on the sea bottom (Fig. 1C). One result would be increased production of non-kelp primary producers, including a higher biomass of understory algae, because the kelp canopy is known to suppress the abundances of understory algal species [12, 25]. The authors also suggest that sea otter-urchin dynamics could have been altered if those conditions led to an increase in invertebrate prey diversity and biomass.

### 1.4 Ecosystem function of *H. gigas* – A new hypothesis

Here we present and test an additional hypothesis of the functional role of *H. gigas*—that sea cow browsing would have increased the biomass of understory algae, and the concentration of drift detritus, thereby increasing the resilience of kelp forests. We define resilience as the capacity of a system, upon being perturbed, to either remain unchanged, or to recover to its state prior to disturbance. The particular case in which a system remains unchanged when perturbed, that is, is maximally resilient, is referred to as resistant. To test the hypothesis we develop a mathematical model that describes the dynamics of a hypothetical kelp forest community off the central California coast, comprising interactions among the giant kelp (*M. pyrifera*), purple sea urchins (*S. purpuratus*), an understory alga favoured by urchins (*Chondrocanthus corymbiferus*), sea otters (*E. lutris*), sunflower sea stars (*P. helianthoides*) and Steller’s sea cow (*H. gigas*) (Fig. 2). We put forth two classes of models, referred to as “Historical” and “Modern”, with the former including the sea cow, and the latter lacking the megaherbivore.

**Figure 2.**
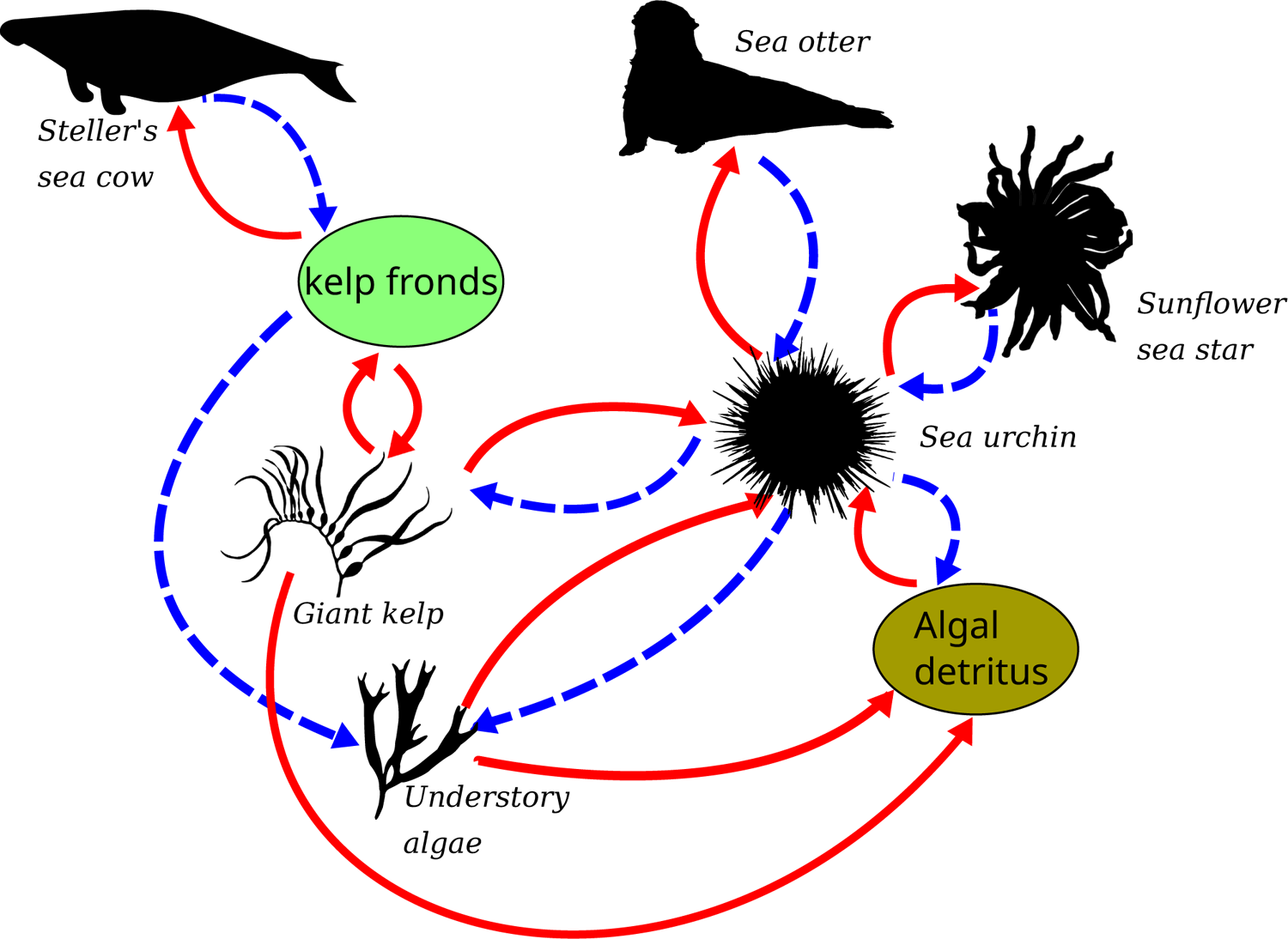

Although kelp density is controlled by numerous factors [2], for the purposes of the model we distill these to dominant factors, including hydrodynamic forces, spatial competition with understory algae, browsing of fronds by Steller’s sea cows, and grazing by purple sea urchins. Extremely high wave energies along the California coast, experienced primarily during winter, are capable of dislodging adult kelp. This phenomenon tends to create bare patches distributed heterogeneously within a forest, regions that are well lit and have substrate available to settlement by kelp, other algae, and sedentary invertebrates [12]. These non-forested patches persist until developing kelp eventually overgrow and shade the area once more. Understory algal density is therefore controlled by spatial competition with kelp on the substrate, light attenuation by the kelp canopy, and sea urchin grazing. Sea urchin density is controlled by sea otter and sunflower sea star predation, and the availability of drift detritus from kelp and understory algae. The urchin population is divided at any given time between sedentary urchins sheltered in substrate crevices, and exposed urchins that are actively foraging. The relative fractions of sedentary versus exposed urchins is controlled by total urchin population size, as well as the availability of drift detritus and the presence of predators [26]. Browsing by sea cows is added to this description of the natural system to build the Historical model.

## 2 Methods

### 2.1 Model system

The models of a modern or historical central Californian kelp forest are systems of biotic interactions (Fig. 2) and physical parameters. Parameters are values that are fixed, or whose variation is independent of other model components, for example incident sunlight or storm events. Variables are partially or fully determined by interactions or dependence on parameters or other variables, for example species populations sizes or densities. The effect of any model parameter or variable is positive if its action on a species or other variable increases the population or numeric size of the other species or variable; negative impacts generate the opposite—decreased population sizes. For example, kelp and algae have a positive effect on the quantity of algal detritus because they produce it, whereas urchins via consumption have a negative effect. The goals of the models developed here are to describe ecological interactions in a manner that results in a feasible forest state when the system is unperturbed, that is species co-exist indefinitely, and to predict system changes during and after perturbation. The following sections outline a mathematical system that describes all the interactions outlined for each model community. Model communities are described with a system of coupled ordinary differential equations, which are simulated and solved using the Julia language. Code used to conduct simulations may be found at the Open Science Framework, URL osf.io/zvnw9. The modeling of physical parameters include ocean temperature, day length and hydrodynamic forces, and is outlined in Supplemental Material. Parameterization of biological parameters is also explained in Supplemental Material.

#### 2.1.1 Species dynamics and biotic interactions

##### Kelp

*Macrocystis pyrifera* population dynamics are modeled as a function of an intrinsic rate of population increase, and predation by urchins and sea cows (when present). Population growth is dependent on both temperature and spatial competition with algae, where kelp are assumed to be the superior competitor because of their greater access to light. Population growth rate also accounts for a steady rate of extrinsic juvenile recruitment, but recruitment is dependent on spatial competition with adult kelp and understory algae, as well as available light at the substrate surface. Urchin predation is a source of direct kelp mortality, whereas sea cow browsing of fronds affects individual productivity and hence the population growth rate, but not mortality. Finally, storm surges are a seasonally stochastic source of mortality.

The intrinsic rate of increase is a daily rate, and should be imagined as a reasonable number of replacements when integrated over a year. For example, *r* = 0.05 would result in a replacement of one adult kelp by *≈* 384 individuals, if growth was uninhibited and exponential. That value is modified by setting *T*_opt_ = 15^°^C. A mean canopy frond density of 20 fronds per individual adult is assumed based on empirical surveys [27]. It is further assumed in the model that kelp density would be twice carrying capacity in the absence of spatial understory competitors. Therefore, if the dependence of the intrinsic rate of increase on temperature and day length is formulated as

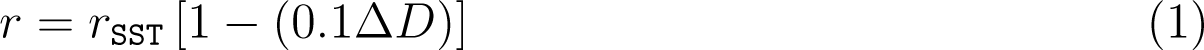

where *r*_SST_ (SST = sea surface temperature) is calculated according to Equation S1 (Supplementary Material) and day length Δ*D* is less than 10 hours, then the population will grow logistically in the absence of predation.

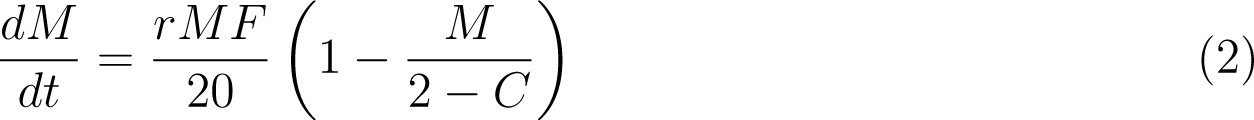

where *M* is kelp density, *F* is frond density and *C* is the density of understory algae.

Urchin browsing of adult kelp is represented as

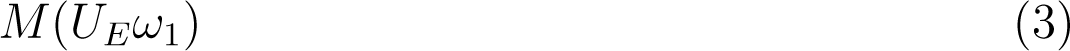

where *U_E_* is the proportion of the urchin population comprising mobile, active foragers (in contrast to sheltered detritivores), and *ω*_1_ is a predator-prey interaction. *ω*_1_ is a functional interaction between urchins and kelp, meaning that the interaction itself is dynamic, changing with prey and predator densities. Here we use an Arditi-Ginzburg-Contois (AGC) function [28], which recognizes prey:predator ratio and predator density as controlling factors in the interaction.

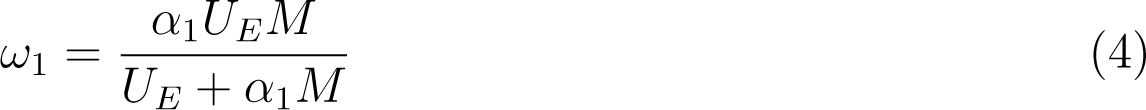

where *α* is the interaction strength between prey and predator (see below).

Local kelp extinction is offset by external recruitment of kelp larvae, which is additional to reproduction by community members. As with the intrinsic rate of increase, the success of recruits is dependent on water temperature, successful settlement and growth, and hence the density of canopy fronds and the amount of light that reaches the benthos. This term is expressed as

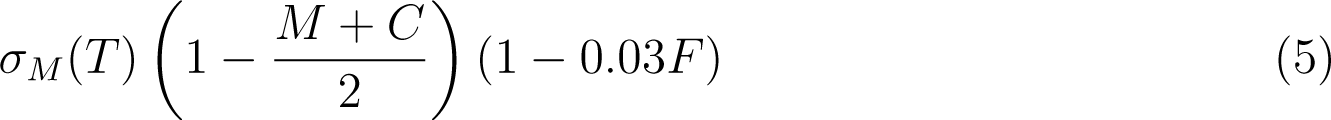

where *σ_M_* (*T*) is the external recruitment rate, *M* and *C* are the densities of kelp and understory algae respectively which dominate the space available for settlement, and 0.03*F* is light availability. The recruitment rate itself is temperature-dependent, expressed as

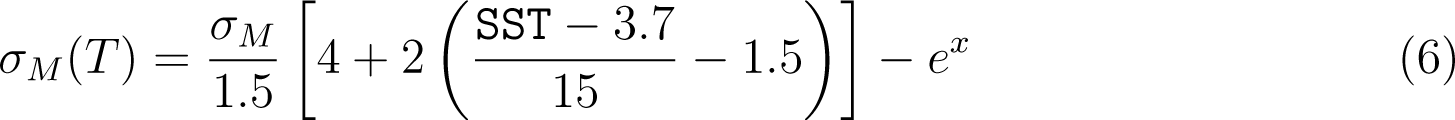

where

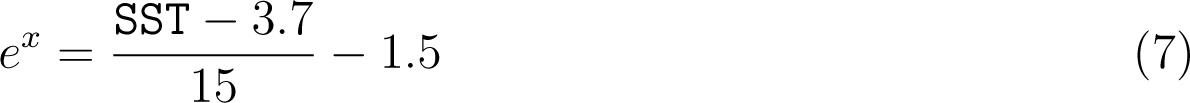

This is a Linex function as employed in Eq. S1.

Combining all these factors, kelp dynamics is therefore expressed as

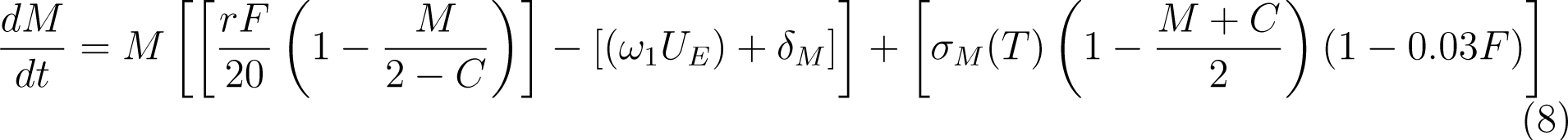

Note that the equation has a discontinuity when *C* = 2, that is, the substrate is completely taken over by understory algae. This is a mathematical aberration only, as the situation implies trivially that the kelp population is then zero, and cannot rise above zero unless new kelp are recruited, and space on the substrate is made available somehow.

##### Kelp canopy

The canopy of kelp fronds at the water’s surface plays several important roles in the community. The canopy absorbs a majority of incoming sunlight and inhibits primary production in the water column and benthos, including that of understory algae. Fronds are one of the major sources of algal macro-detritus, hence a food resource for urchins, and they were apparently the sole food for *H. gigas*.

Frond density is estimated in the model as 20 per *M. pyrifera* individual adult kelp [27]. Fronds are consumed by sea cows when they are present in the system, which in turn creates a negative feedback to kelp growth rate (*dM/dt*). Fronds also have a natural senescence rate at which they age and detach from the stalk. Thus,

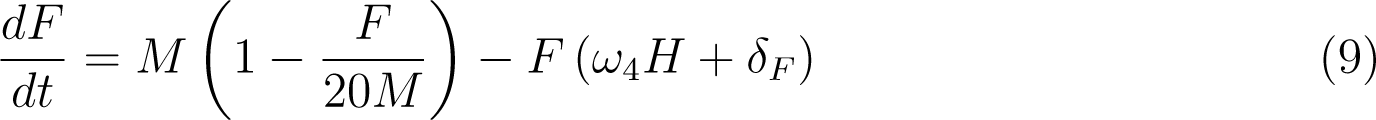

where *F* is frond density, *H* is sea cow density, *ω*_4_ is the interaction between fronds and sea cows of the AGC form (see Eq. 4), and *δ_F_* is the frond senescence rate.

##### Drift detritus

Both kelp and the understory algae in the model contribute to drifting macrodetritus. Detritus is produced as a fixed fraction of frond and understory algal density, and becomes refractory (inedible) also at a fixed rate. Thus,

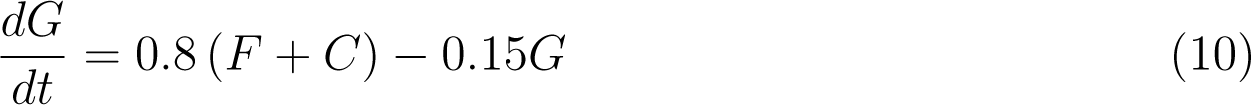

where *G* is detrital abundance, 0.8 is derived from empirical measures of detritial production by another kelp species, *Laminaria hyperborea* [29], and 0.15d*^−^*^1^ is the refraction rate.

##### Understory algae

This group is represented in the model by the red alga *Chondrocanthus corymbiferus*, which has been shown to be nutritious for urchins, readily consumed when available, and capable of sustaining urchins after kelp density has been reduced by large storm events [30]. The alga contributes to the detrital pool, competes spatially with kelp on the substrate, is light-inhibited by the kelp, and is grazed by actively foraging urchins. Like kelp, its rate of intrinsic increase is temperature-dependent, with *T*_opt_ set here at 13^°^C, slightly cooler than that of the kelp, based on the more limited southern extent of its range. Dynamics are given by

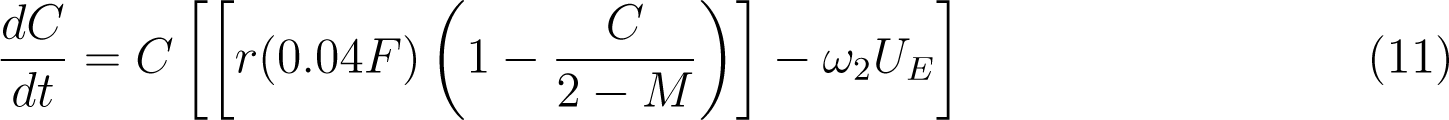

where (0.04*F*) is the degree to which population growth is impacted by kelp frond density. That is, when the kelp canopy reaches maximum density, we assume that growth of *C. corymbiferus* is inhibited by 96%, a value slightly larger than the average 90% of light attenuation in healthy forests [12]. *ω*_2_ is the predation rate by exposed, actively foraging urchins.

##### Sea urchins

Two common species of strongylocentrotid urchins exist in kelp forests of the Pacific northwest: *Strongylocentrotus purpuratus* and *Mesocentrotus franciscanus*, the purple and red sea urchins, respectively. Both are treated as a single unit in the model, but in reality, although the two species are associated with the transition between forested and barren system states, the purple urchin is by far the dominant species in barrens. This is most likely due to this species’ greater ability to withstand a poor nutritional environment. The major controls of sea urchin dynamics are population density and food supply.

Urchins prefer to remain sedentary and sheltered from predation using the topography of the benthic substrate. When sheltered and sedentary, urchins feed primarily on drifting algal detritus. All such detritus in the model is produced by *M pyrifera* and *C. corymbiferus* (Eq. 10). Nevertheless, a small fraction of the population can usually be found exposed and mobile even when detrital production is high [31]. Mobile urchins consume both detritus and living algae, and are responsible for the transition from the forest to the barrens state. The proportion of the urchin population that is actively foraging increases as population density increases, possibly because of crowding in the substrate. It also grows if detrital concentration declines [26], and competition within the sedentary lifestyle increases. There is thus a positive feedback between active foraging and the necessity of leaving crevices.

The proportion of exposed urchins, *U_E_*, is modeled as follows. First it is assumed that there is a finite amount of space available in the substrate for sheltering urchins. Using data available for Monterey Bay, California [31], we estimated that if total urchin density is less than the amount of available space, then the proportion of exposed urchins, that is, mobile foragers, may be estimated as

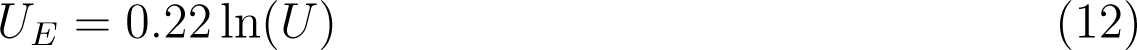

(*r*^2^ = 0.45, *p <* 0.0001). There is also a weaker dependency on kelp density, as indicated by the gonadal condition (gonad index) of the urchins. We assumed detrital density to be a direct function of adult urchin density as measured with those data, and that near the carrying capacity of sheltered urchins the proportion of exposed urchins is approximately 0.35 [31]. Thus, we modeled the relationship between exposed urchins, total urchin density and available drift detritus as (multiple least squares regression; *R*^2^ = 0.45, p*<*0.0001) where *U* is total urchin density and *G* is detrital density (Fig. 3).

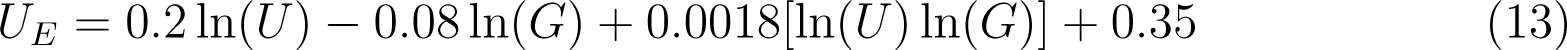

**Figure 3.**
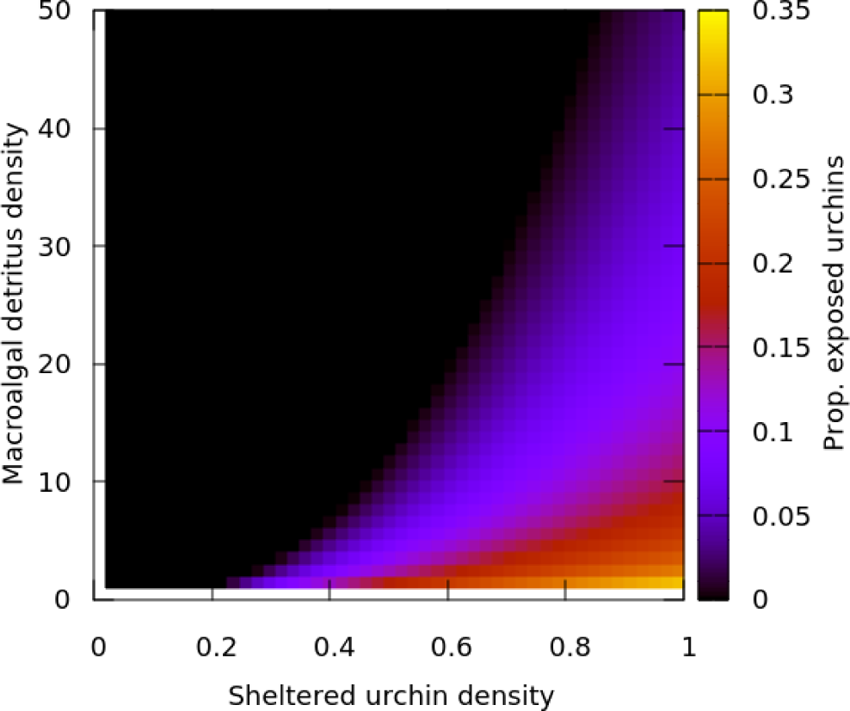

Growth of the total urchin population is then determined by the foraging activity of exposed urchins, detritivory by both sheltered and exposed urchins, and predation by both major predators of the urchins: sea stars and sea otters.

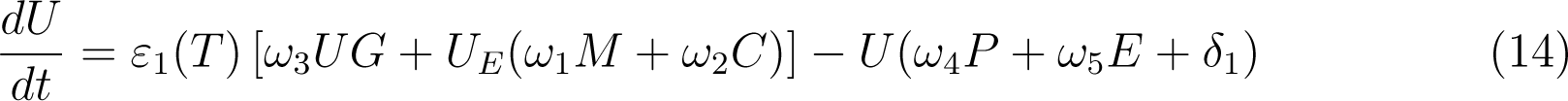

where *ε*_1_ is a temperature-dependent ecological efficiency of the urchins of the same form as Equation S1, *ω_n_* are predator-prey coefficients, *P* is sea star density, *E* is sea otter density, and *δ*_1_ is the urchin mortality rate.

##### Sunflower stars

*Pycnopodia helianthoides*, like the urchins, is assumed to have SST-dependent rates of prey interaction and ecological efficiency of the same form as Equation S1. Predation of sheltered versus exposed urchins are tracked separately, and the interaction between sunflower sea stars and urchins is therefore expressed as

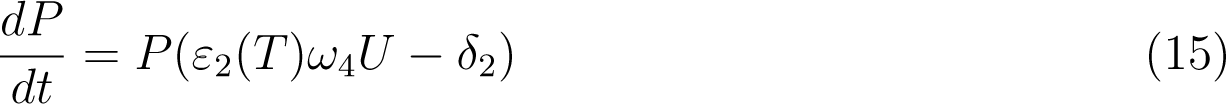

where again, *ε*_2_ is the ecological efficiency, *ω*_4_ is the predator-prey interaction term, and *δ*_2_ is a mortality rate.

##### Steller’s sea cow

*Hydrodamalis gigas* was apparently an obligate canopy feeder, based on observations of its feeding and its bouyancy. No instances of non-human predation were ever observed, although anecdotal behavioural reports suggest that the animals were wary of orcas and swam in formations that may have shielded juveniles [32]. It is also reasonable to speculate that large sharks, particularly great whites (*Carcharodon carcharias*), may have preyed on juveniles. Nevertheless, no predation is included in the model. *H. gigas* dynamics were therefore controlled by frond availability and a natural mortality rate.

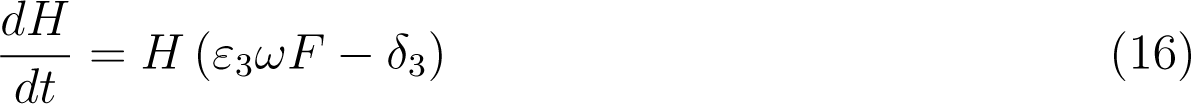

where *ε*_3_ is the ecological efficiency with which consumed kelp was “converted” to new sea cow individuals. *δ*_3_ is the natural mortality rate.

##### Sea otters

Sea otters, *Enhyda lutris*, are present only when kelp are sufficiently abundant, and urchins are therefore healthy. Otters reject urchins with low gonadal and other soft tissue content, generally avoiding foraging in urchin barrens. Otter population dynamics are therefore ultimately kelp-dependent. Given the ephemerality of sea otters in our model community, we assumed that the otters are drawn from a fixed metapopulation pool. Further assuming that urchin gonadal index is a function of kelp density, and thus that the presence of otters preying on urchins is likewise dependent on kelp density, we used published data of the relationship between gonadal index and otter predation [31] to derive a logistic relationship between kelp density and the probability that otters are present and foraging (Supplementary Material).

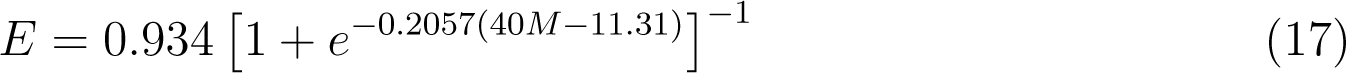

where *E* is sea otter density in the community, and *M* is kelp density.

### 2.2 Perturbation

Model communities were subjected to four types of perturbation: no perturbation; a MHW to simulate the warm Pacific Ocean blob; an outbreak of SSWD; and coincident warming and disease. Model communities were constructed as described above and simulated for a burn-in period of 200,000 days, at which point the state of the community was described as forest, or quasiperiodic, this latter description capturing oscillations between the forest and barren states (see Results). Perturbations were then applied to the models.

In the case of no perturbation, the model was simulated for an additional 14600 days (40 years) post burn-in. For each of the perturbed models, perturbations were initiated precisely 10 years after the burn-in (day 203,650). The MHW, or blob perturbation, consisted of an abrupt increase of temperature 3.0^°^C in excess of the seasonal cycle, and lasted for 2 years (730 days), after which temperatures returned to the seasonal cycle. The onset of SSWD was also initiated 10 years after burn-in, and was implemented as an exponential decline of the *P. helianthoides* population, modeled from empirical data of population decline from central and northern California [19].

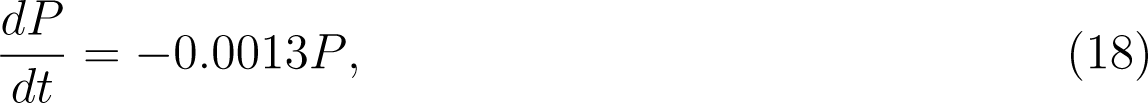

after which the sea star population was allowed to recover. Models were therefore simulated for 30 and 37 years after the termination of disease and warming respectively.

A total of 100 simulations were performed for each of 16 total model-perturbation combinations. This included the Historical and Modern models (i.e. with and without sea cows) with either no perturbation, warming, disease, or warming plus disease, each with and without sea otters. This resulted in a total of 1,600 simulations. Each simulation consisted of the solutions to the appropriate system of differential equations as outlined in the previous sections.

### 2.3 Simulation analysis

#### 2.3.1 Unperturbed simulations

Unperturbed simulations were conducted to measure the frequencies at which various states occur within ranges of the model parameters, without the impact of external factors. Simulation outcomes were classified according to the relative abundances of system species: simulations in which giant kelp are dominant and remain abundant represent the forest state, and those in which urchins dominate numerically and kelp abundance is very low were classified as transient barrens.

#### 2.3.2 Perturbed simulations

Simulations, perturbed as described above, were classified according to the scheme outlined for unperturbed simulations. In this case, however, only simulations that were in the forest state at the end of the burn-in period were analyzed, as we are interested in the impact of the perturbations on forests. Those simulations could then either remain in the forest state after initiation of a perturbation, or transition to a barrens. The relative frequency of each type of response was recorded for each model-perturbation combination, of which there are 12; there are two model classes, Historical and Modern, and within each there are two sub-classes, systems with sea otter predation and systems without. Furthermore, each subclass was subjected to three different perturbations: ocean warming, sea star wasting disease, and a combination of warming and disease.

#### 2.3.3 Measuring resilience

Resilient simulations are those that remain in the forest state after perturbation, or by definition here, return to a forest state within ten years after the end of a perturbation. The frequency of resilient simulations were recorded for each set of simulations of a model-perturbation combination, along with the corresponding frequency of simulations that transitioned permanently away from the forest state.

Resilient simulations were further sub-categorized according to whether they exhibited no state transitions, or underwent one or more transitions but subsequently recovered permanently (within the time frame of the simulation) to the forest state. Simulations that did not undergo any transitions were categorized as resistant. The resilience of non-resistant, but recovered simulations was described with the time of onset of the first transition to barrens, and the time of the last transition back to forest. These two measures encompass one classic definition of ecological resilience, which is recovery time [33]

## 3 Results

### 3.1 Community states

Three states emerge from the unperturbed models, two of which have dense kelp forests. All species persist in the first forest state, corresponding to the conventional concept of a healthy kelp forest (Fig. 4A); we refer to this state hereon as simply forest. The other forested state is one in which *S. purpuratus* and *P. helianthoides* fail to persist, or do so at densities less than 1% relative to their initial values. We term this a “low diversity forest”. The third state is a quasiperiodic oscillation between healthy forest and urchin barrens, the latter condition being one in which algal densities are drastically reduced (Fig. 4B). The barrens state is, however, unstable in our model because of continuous temperature-dependent recruitment of kelp and understory algae from external sources. Therefore, healthy forest may recover after transition to barrens, although the timescale varied among simulations.

**Figure 4.**
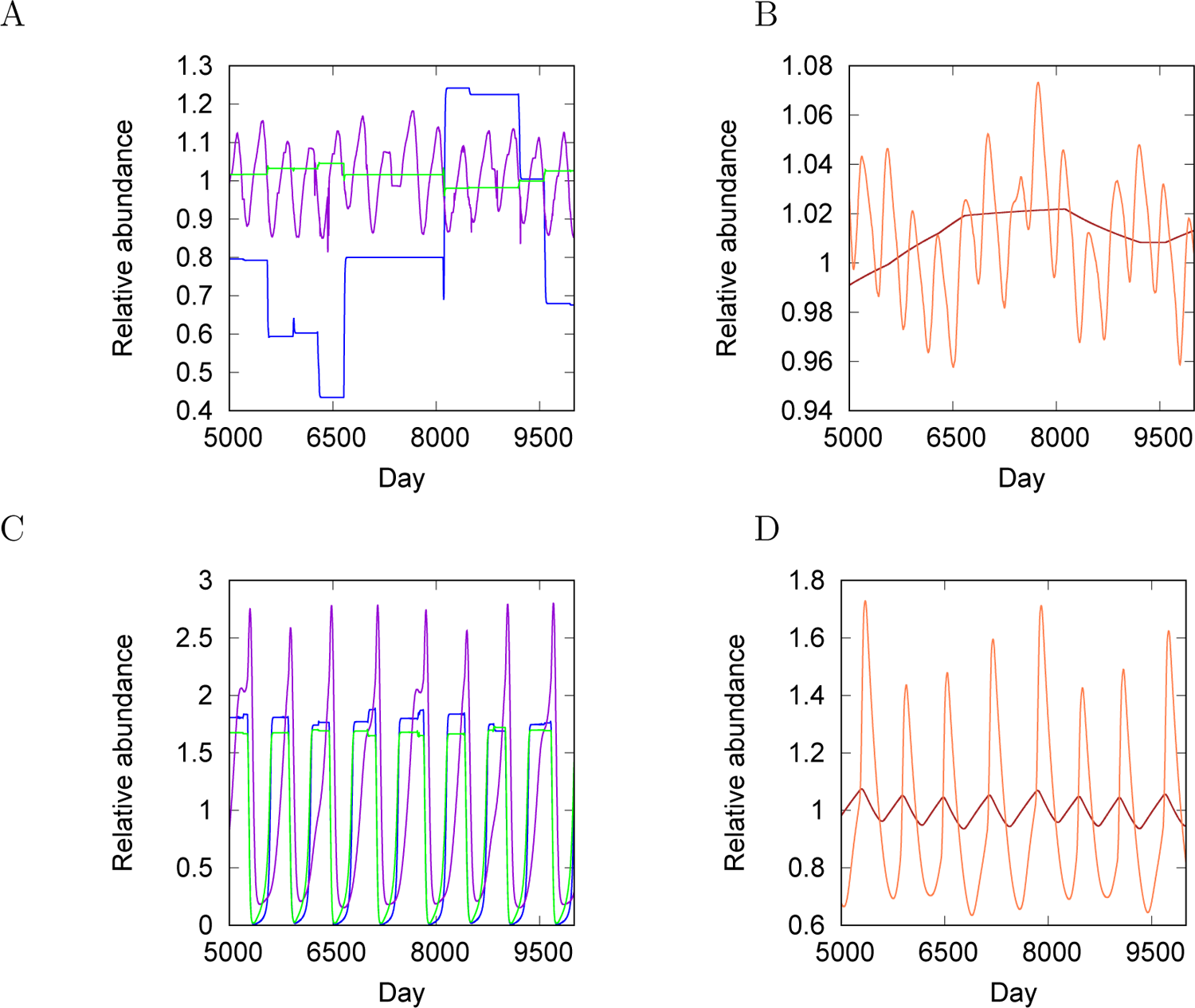

One hundred simulations of each model type showed statistically indistinguishable frequencies of the three states among the models; forest, low diversity forest, and quasiperiodic oscillation (chi-squared test, *χ*^2^ = 8.9215, p=0.1780). The ensemble of simulations across all model types yielded frequencies of 66.5% forest, 25.5% quasiperiodic oscillation, and 7.7% low diversity forest. Given that systems exhibiting quasiperiodic oscillation also exhibit intervals of forest, and that intervals spent in the forest condition during oscillations are at least as lengthy as barrens intervals, then under unperturbed conditions, areas with the potential to host *M. pyrifera* should be forested in least 74.4% of occurrences. This is consistent with the observed widespread occurrence of giant kelp forests.

Sea urchin ecological efficiency differs significantly among the states (Fig. **??**) and predicts when an unperturbed simulation will produce either the forest or quasiperiodic state (Supplemental Table S2). The parameter is estimated as having a mean value of 10% (Supplemental Material), but varies among simulations with the addition of a small error parameter drawn from a normal distribution of mean zero and standard deviation of 0.003. This is reflected by the significantly different distributions of the parameter among the states (ANOVA, F=55.32, p*<*0.0001, Scheffe’s multicomparison test) (Fig. **??**A). The dominance of urchins in the system increases with ecological efficiency, resulting in the three alternative states.

Considering the two focal states of this study (healthy forest and quasiperiodic) and the difference between the distributions of urchin ecological efficiency between those states, we evaluated the dependence of transitions between states on ecological efficiency with a logistic regression (*χ*^2^ = 47.65, *R*^2^ = 0.441, p*<*0.00001) (Fig. **??**B). The emergence of the quasiperiodic state begins at an efficiency of approximately 0.102, with an increasing probability of emergence as efficiency increases. The healthy forest state is still possible at those values of efficiency though, and the system is therefore capable in being in one of the two alternative states at higher efficiency values.

#### 3.1.1 Forest types

As this study is focused on the response of the forest state to perturbation, it was necessary to establish whether the forest state itself is identical under the different models. A comparison of population sizes of species common to all the models (*M. pyrifer*, *C. corymbiferus*, *S. purpuratus* and *P. helianthoides*) across unperturbed models, during the final year of each simulation shows that the forest state differs significantly among the models (MANOVA, Wilks’ *λ* = 0.4043, *F* = 9286.86, *p <* 0.00001). The differences are driven by an inverse relationship between giant kelp and understory algal abundances (Fig. 6) (Supplementary Figure 2). Models or systems without sea otter predation have greater abundances of *M. pyrifera* relative to *C. corymbiferus*. The Historical model with sea otters has the lowest relative abundance of giant kelp, whereas the Historical model without sea otters has the highest relative abundance. Thus the forest state across all models is itself heterogeneous.

**Figure 5.**
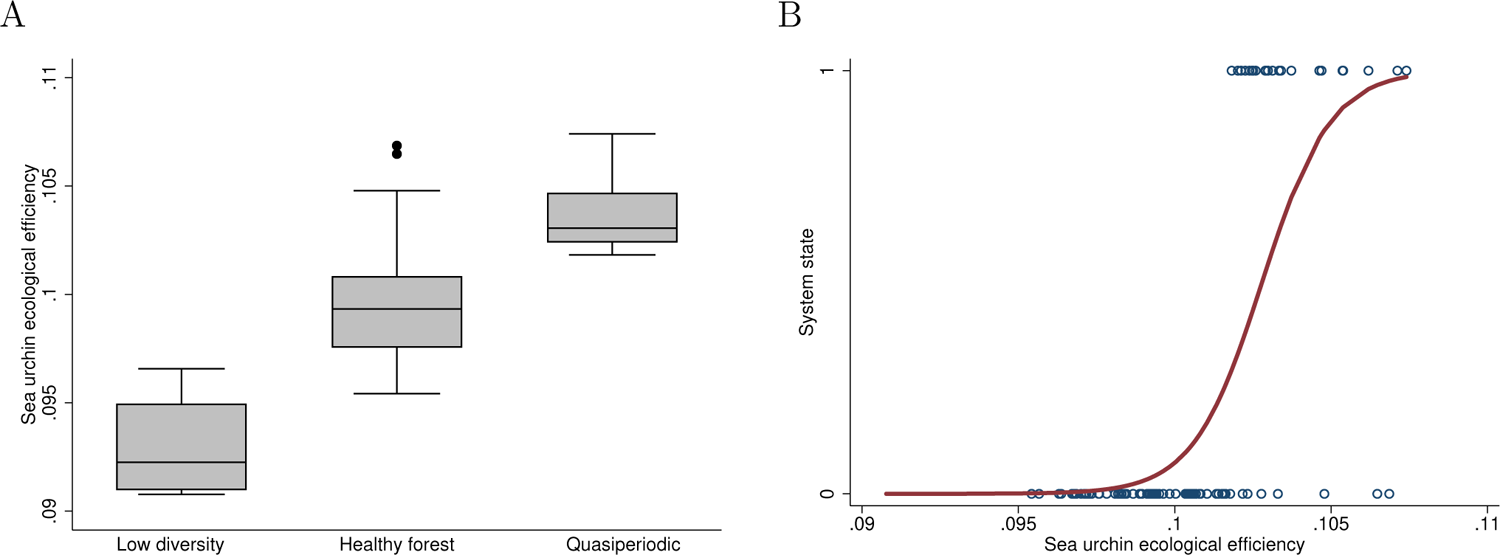

**Figure 6.**
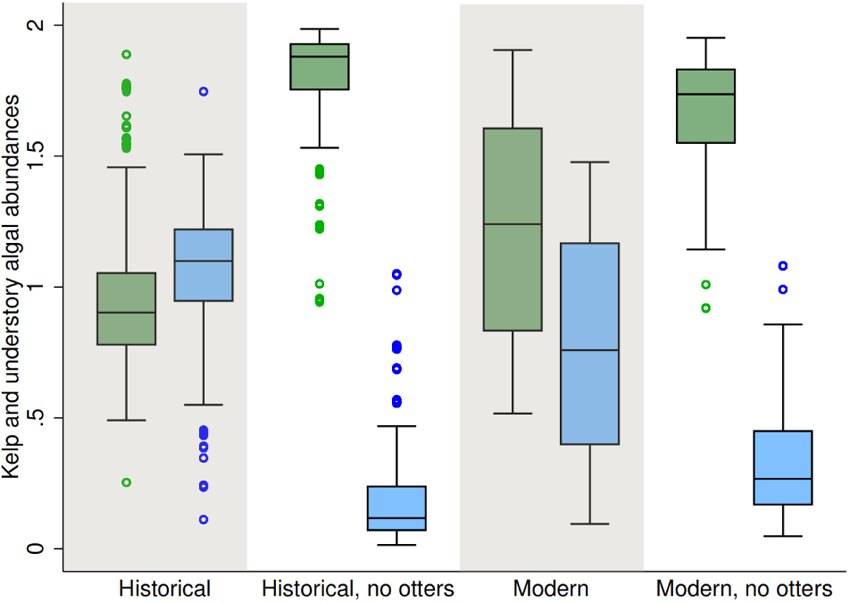

### 3.2 Response to perturbation

#### 3.2.1 State transitions

Responses to perturbation were classified as either being in a stable forest state at the end of the simulation (Fig. 7A,B), or having transitioned to a quasiperiodic state (Fig. 7C, D). The frequencies of transitions were compared pairwise among model-perturbation combinations. Recalling from the previous section that model types did not differ in the frequency with which they produce the forest state when unperturbed, we expect that any differences when perturbed are caused by perturbations only, and do not reflect stochastic differences among simulations.

**Figure 7.**
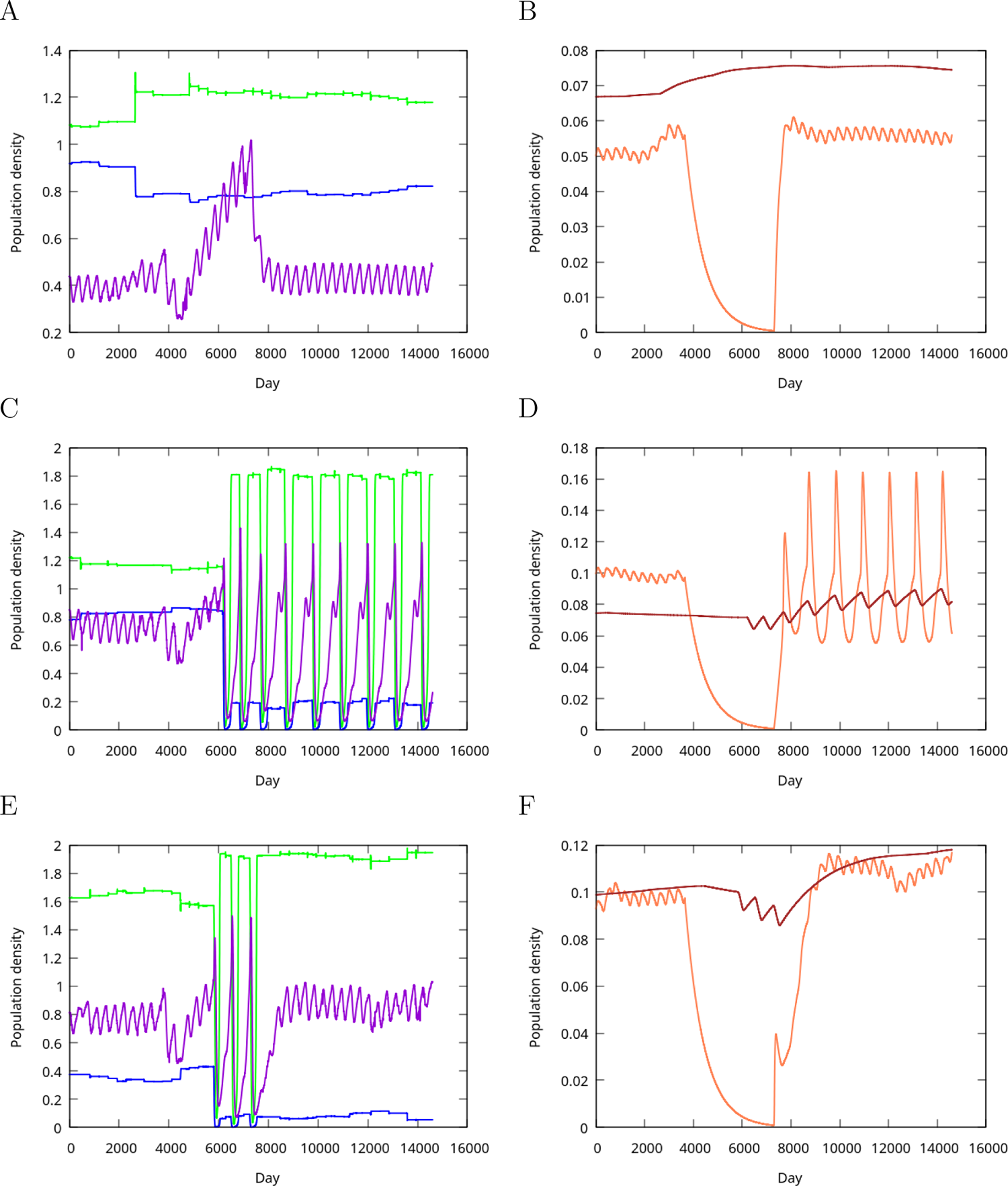

Almost all model-perturbation combinations exhibited similar frequencies of transition to the quasiperiodic state, with the exception of the Historical model when it was subjected to a combination of MHW and SSWD. This model produced a significantly greater transition frequency when compared to other models (*χ*^2^ comparisons, *α* = 0.05. The pooled frequency of all other model-perturbation combinations is 3.29%, whereas it differs significantly at 32.23% when warming and disease are applied to the Historical model (*χ*^2^ = 36.718, p*<*0.00001).

#### 3.2.2 Resilience and resistance

Resilient simulations were those that began in the forest state prior to perturbation and ended in the forest state. Simulations were classified as resistant (maximally resilient), if barrens never appeared (Fig. 7A, B), or less resilient if barrens occurred but the system subsequently returned to stable forest (Fig. 7D, E). Resistant systems exhibited temporary increases of sea urchin density without state transition because of either a relatively more favourable tolerance to warming compared to their sunflower sea star predators, or because of release from predation as disease reduced sunflower sea star density, or both. In contrast, less resilient systems always underwent multiple state switches between forest and barrens before returning permanently to the forest state.

The relative frequencies of resistant and resilient simulations varied significantly among model types and perturbations (Supplementary Material), and model scenarios fall into several groups. All simulations that ended in the forest state were either resistant or less resilient (Fig. 8) for each model-perturbation combination. There is an inverse relationship between the frequencies, that is, a larger number of resistant simulations must necessarily mean a smaller number of less resilient ones for each model-perturbation combination, and *vice versa*.

**Figure 8.**
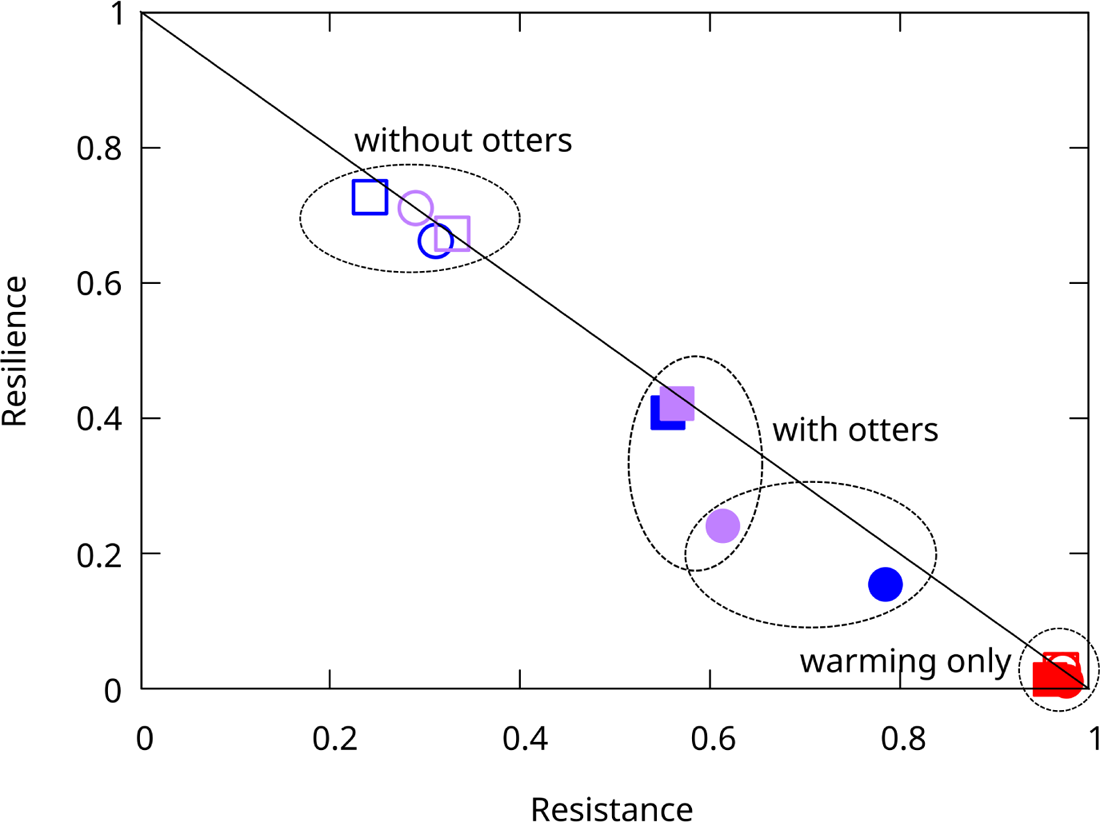

The group with the greatest number of resistant forests were those perturbed by warming only, regardless of model type. In contrast, the group with the fewest resistant simulations, and hence the greatest number of less resilient ones, comprised both the Historical and Modern models without sea otters, when subjected to a combined perturbation of warming and disease. Intermediate were the Historical and Modern models with sea otters, and when subjected to either disease, or the combination of warming and disease. There is no significant difference between the Modern system and the Historical one when the latter is subjected to both warming and disease (compared to Modern disease, and Modern warming plus disease; *χ*^2^ =3.24 and 3.34; *p* =0.072 and 0.067 respectively). The Historical model subjected to disease only, however, is significantly more resistant than the Modern models (compared to Modern disease, and Modern warming plus disease; *χ*^2^ =12.66 and 12.78; *p* =0.0004 and 0.0003, respectively).

#### 3.2.3 Onset and recovery from transitions

Resilient simulations were characterized by comparing the times of onset of the first transition to barrens, and the time of permanent recovery to the forest state. Models perturbed by warming only were excluded from the analysis. There were no significant differences of either the time of onset or recovery among those models perturbed by SSWD only (ANOVA, n=144; onset, F=1.06, p=0.368; recovery, F=0.68, p=0.567), or SSWD plus MHW only (ANOVA, n=131; onset, F=0.70, p=0.5; recovery, F=0.88, p=0.418), nor did models differ in recovery time if the perturbations were pooled (ANOVA, n=293; recovery, F=0.61, p=0.747). Onset time differed among the pooled Historical models (ANOVA, n=293; onset, F=2.06, p=0.048), with the differences being explained by earlier onset times when sea otters are absent from the system (Supplementary Figure 2). This is also true for the Modern model only when the perturbation is the combination of warming and SSWD. In other words, sea otters delay the onset of any transitions to urchin barrens when Steller’s sea cow is present. They fail to play a similar role in the Modern system when perturbed by SSW only.

Historical systems with Steller’s sea cow and sea otters present had a mean onset time *∼*7.5 years after the onset of perturbations, whereas the absence of the sea otter resulted in an earlier mean (*∼*5.9 years). Modern systems also differed based on the presence or absence of the sea otter, in addition to whether the perturbation was SSWD only, or the combination of warming and disease. The presence of the sea otter made no difference to onset time if the system was perturbed by SSWD only, with a mean onset time of 2,281 days (*∼*6.25 years). When subjected to both SSWD and warming, however, the Modern system with sea otters had a mean onset time of *∼*7.35 years, whereas without sea otters the mean onset time was *∼*6.45 years.

### 3.3 Predicting transition

Variation of the pre-perturbation conditions might determine whether they are resistant or resilient, potentially serving as predictive indicators in Modern systems of how they would respond to perturbation. We tested this with a principal components analysis (PCA) of population sizes of the four species common to all model or system types: *M. pyrifera, C. corymbiferus, S. purpuratus* and *P. helianthoides*. We restricted the analysis to population sizes during the final year before the onset of perturbation, where the perturbations were either SSWD, or SSWD plus MHW. The PCA summary of the state of each simulation prior to perturbation differentiates between resistant and resilient simulations (Fig. 9A). Loadings of the variables indicate that the relative abundances of *M. pyrifera* and *C. corymbiferus* are the most important determinants of whether a response will be resistant or resilient (Supplemental Tables 3-4).

**Figure 9.**
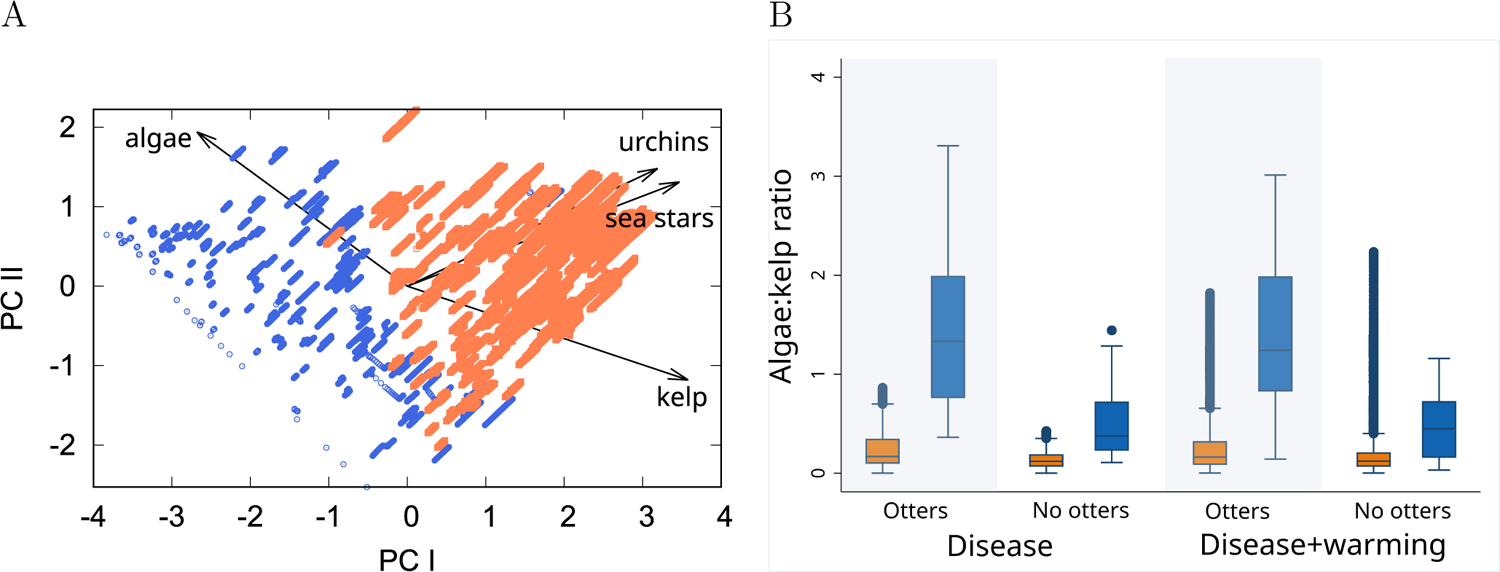

We summarized this by comparing the ratios of understory algae to kelp population sizes during the year before perturbation, demonstrating that those ratios differ significantly between resistant and resilient responses among all the Modern model types (Fig. 9B). In each model type, that is with or without sea otters, and subjected to either type of perturbation, systems that were subsequently resistant to any transient transitions to quasiperiodicity were those with significantly greater ratios of understory algae to kelp. Furthermore, understory algae have larger population sizes in the presence of sea otters, presumably because their predation of sea urchins releases the algae from urchin predation, allowing them to compete more successfully against kelp.

## 4 Discussion

### 4.1 Alternative states without perturbation

Unperturbed simulations of all model types (Historical, Historical without otters, Modern, and Modern without otters) revealed three possible alternative states of *M. pyrifera* giant kelp communities (Fig. 4), termed here “low diversity forest”, “healthy forest” (or simply forest) and “quasiperiodic”, the latter of which transitions frequently between a forested community and urchin barrens. All are intrinsic features of the system, and need not be a response to the external environment. Cyclical variations do occur between the coupled predator-prey echinoderms, but those are driven by annual seasonal temperature variation (Fig. 4A, B) that affects the species physiologically. Quasiperiodic oscillation, in contrast, is of a lower frequency, with wavelengths exceeding a year (Fig. 4C, D), and therefore driven, at least partly, by intrinsic system dynamics.

All model types generate these states at indistinguishable frequencies, implying that system state is not dependent on the differences between the models. Why then are the models capable of generating different states if unperturbed? Aside from species composition, the other source of variation within a model is the estimation of several parameters: rates of intrinsic population increase of giant kelp and understory algae (*C. corymbiferus*), and the ecological efficiencies of the consumers *S. purpuratus* and *P. helianthoides*. Sea urchin ecological efficiency emerged as the single explanatory parameter (Supplemental Table S2). This parameter varies to reflect uncertainty and true variation of its value *in vivo*. The generation of three discrete states represents a probabilistic system bifurcation, in which state depends on a noisy parameter [34]. Higher values of urchin efficiency increase the likelihood of quasiperiodic variation and hence urchin barrens. The ecological implication is that any factors leading to an increase of that efficiency, or to its greater influence on population growth, are also likely to give rise to quasiperiodic transitions between kelp forest and urchin barrens. Sea urchin ecological efficiency is temperature dependent (see Eq. **??**). Therefore, ocean warming or cooling toward optimum temperature will increase the probability of quasiperiodicity.

### 4.2 Multiple types of forest

The models also reveal a diversity of forest types, differentiated by the relative densities of giant kelp and understory algae. There is an inverse relationship between kelp and understory algal abundances in three of the model types, with the “Historical with sea otters” model being an exception relative to the others (Fig. 6). The relationship is pronounced in the Historical and Modern models when sea otter predation is absent. In those models, both kelp and understory algae are subject to more intense grazing by foraging sea urchins because of lower predation pressure. The understory algae, however, are also subject to light inhibition by the kelp canopy, and therefore reach their lowest abundances in those systems. Kelp, as a consequence, attain their greatest abundances because of decreased spatial competition with the understory algae on the substrate. The Modern system with sea otters has overlapping densities of kelp and understory algae, although kelp still dominate, because the relative abundances are now controlled primarily by spatial competition, and shading of the understory by the kelp. The relationship is reversed in the Historical model, with the abundances being more nearly equal, because of browsing by Steller’s sea cow. The megaherbivore controls kelp abundance both by inhibiting growth with the removal of kelp fronds, and by allowing the algae to be more competitive due to greater light penetration [35, 36].

The models therefore predict that kelp forests today should vary in the relative abundances of giant kelp and understory algae that are favoured by urchins, and that the variance is dependent on the presence or absence of predatory otters. We are not aware of any prior theoretical or empirical studies that have suggested this variance, but it is amenable to empirical verification. The role of the sea otter is not surprising, but conventional thought is that thir positive impact is the result of their control of sea urchins who then exert less pressure on kelp. Here we suggest an additional route whereby decreased grazing on understory algae by urchins in the presence of sea otters allows the algae to attain greater densities, being limited then solely by shading and competition with kelp. Understory algae thereby become a more important alternative food resource for urchins. Thus, the sea otter as a keystone predator may be important for persistence of the forest state and increased primary producer diversity via both direct and indirect interactions.

#### 4.2.1 A no-analog community

Our results further predict the existence of a novel type of forest that would have existed in the past when Steller’s sea cow was extant. Frond browsing by the sirenian would have had a positive effect on understory algal abundance, further amplified by the sea otter, resulting in kelp forests where the abundance of understory algae could match or even exceed that of the kelp. Today there are no megaherbivores that feed on kelp forest canopies, and herbivory by invertebrates and herbivorous fish are unlikely to open up the canopy in ways that Steller’s sea cow was able to. The decline and extinction of *Hydrodamalis* species would therefore have marked the demise of a type of kelp forest for which there is no analog today. Results presented here suggest that this would have had significant effects on both forest resilience and the probability of transitions to barrens when perturbed by MHW warming and SSWD. The dynamics of this novel forest and the impact of Steller’s sea cow, however, are complicated.

### 4.3 Resilience and transition

Healthy forests, when subjected to either MHW and/or SSWD, were generally, or at least be resilient, returning to the forest state after one or more transient transitions to barrens (Fig. 8A, E). Alternatively, the system could undergo a transition to quasiperiodicity (Fig. 8B), where it permanently alternated between forest and barrens. Most of the model-perturbation combinations were highly resilient to permanent transitions to the quasiperiodicity (96.71% of simulations). One should therefore expect that a majority of kelp forests subjected to MHW and/or SSWD will not undergo long-term (10 or more years) transitions to an unstable, alternating state. The Historical model, with sea otters present, is a significant exception to this prediction, with our results predicting a collapse of those forests more than 30% of the time when subjected to a combined perturbation of MHW and disease. This is notable, and worth further exploration.

Several parameters describing the ecologies of kelp, understory algae, and the invertebrate echinoderms, are temperature sensitive in the model and decline as temperature increases. These include the intrinsic rates of population increase of the producers, and the predatorprey interaction coefficients and ecological efficiencies of consumers. For example, increases of water temperature decreases *P. helianthoides* growth rates [37] Furthermore, SSWD reduces the interaction between sea urchins and sunflower sea stars from the system, and rapid declines of *P. helianthoides* have been documented to lead to increases of the fraction of urchin populations that are exposed and foraging, and declines of the urchins’ nutritional state [31]. In the Modern system, urchin populations would then be controlled by sea otters only, and this is apparently sufficient to maintain the system in a forested state. In the Historical system, however, Steller’s sea cow continues to exert a negative impact on the kelp both through consumption of fronds, and subsequent release of understory algae from shading. The combined effects of urchin release from sea star predation, depression of kelp growth rates, and sustained sea cow grazing, is sufficient to tip the system into the quasiperiodic state.

The Historic system when subjected to SSWD only does not have a high probability of transition because the kelp rate of intrinsic increase is not depressed by warming. Paradoxically, when the system lacks sea otter predation, it is also highly resistant to the combined warming-disease perturbation, which would appear to be inconsistent with the above explanation because urchins are now under no predation pressure. We propose a type of intermediate disturbance hypothesis (IDH) [38] to explain these results. The classic IDH seeks to explain the maximization of biodiversity at intermediate levels of disturbance by a minimization of competition. Here we propose that the probability of transition to quasiperiodicity of the Historical system is *maximized* at an intermediate level of predation. The lowest level of disturbance occurs when sea otters are present and the system is perturbed by disease only; resistance is maximized because kelp and understory algae are not affected by the disease (Fig. 10). The intermediate situation arises when they are subjected to MHW and SSWD, and urchins are released from sea star predation but not predation by sea otters. Maximum disturbance occurs under warming plus disease when sea otters are absent. In the maximum disturbance case, we propose that the urchin population is so tightly coupled to the dynamics of its prey (kelp and understory algae) prior to any disturbance because otters are absent, that warming weakens this producer-herbivore coupling and the impact of the urchins on their prey sufficiently to prevent the system from transitioning into quasiperiodicity.

**Figure 10.**
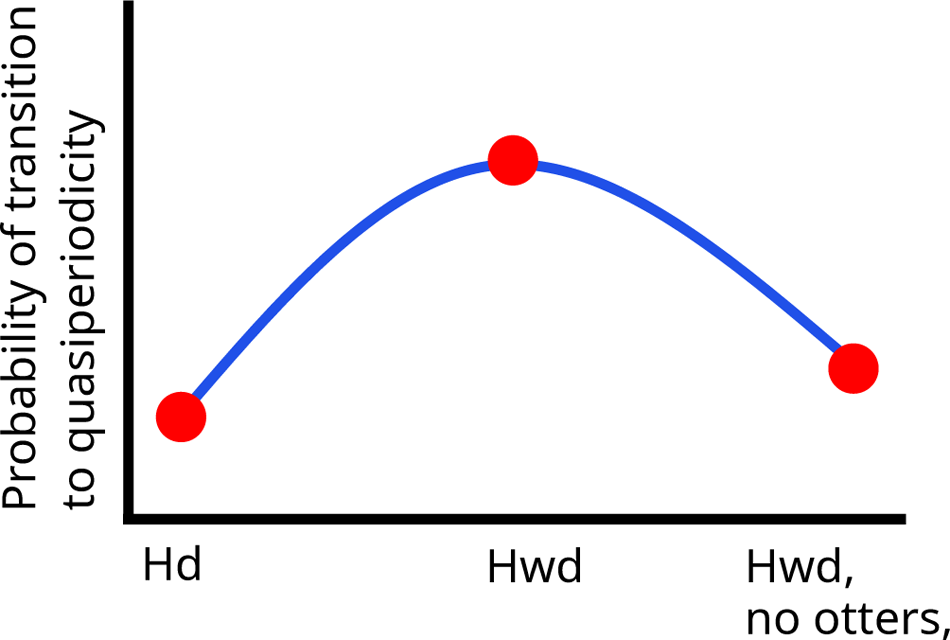

**Figure 11.**
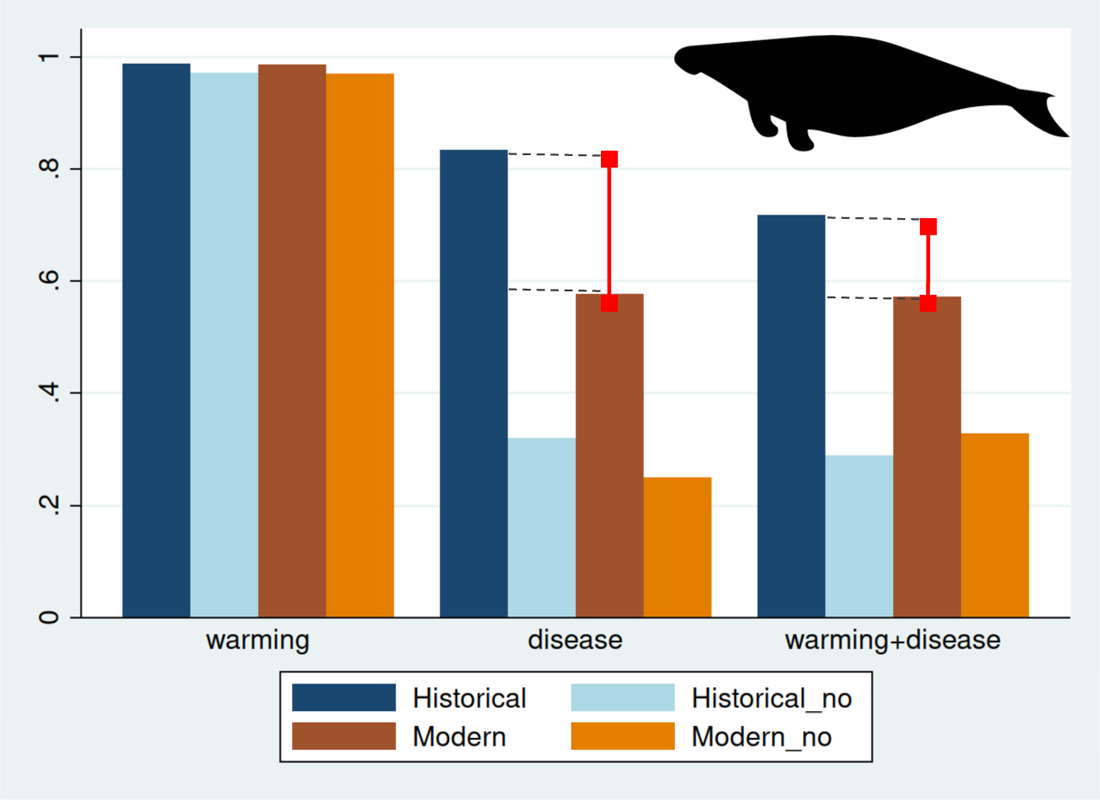

The complicated nature of the relationship between perturbations and transitions proposed here is typical of ecological systems of even modest diversity and complexity, and understanding the combined effects of stressors can be challenging [2]. Filbee-Dexter and Scheibling [7] have suggested that the transitions between forest and barrens may have different thresholds and alternative state dynamics, typical of the hysteresis observed and theorized to be commonly associated with state transitions [39]. Rogers-Bennett [40] has emphasized that the conventional predator-driven deforestation paradigm may be too straightforward, overlooking the roles of other system feedbacks.

#### 4.3.1 Transitions and extinction

The susceptibility of the Historical system to transition from the forest to the quasiperiodic state when subjected to MHW plus SSWD provides an alternative hypothesis to explain the numerical and geographic contraction of the sea cow. Current hypotheses postulate either Indigenous hunting or glacial disruptions as explanations of why Steller’s sea cow was demographically and geographically limited by the time that it was first described scientifically [41, 42]. Here we suggest that warming temperatures since the end of the last glacial termination increased the likelihood and frequency of both MHW and outbreaks of diseases that affect urchin predators, and hence the frequency at which urchin barrens occur. Local transitions of kelp forests to quasiperiodic or barrens states for decadal durations would have been unfavourable to the large, slow reproducing mammal, thereby increasing rates of local extirpation. Historical restriction to the most poleward parts of the species distribution, as observed in the fossil and historical records [23, 41], would be consistent with this hypothesis.

### 4.4 Degrees of resilience

Resilient simulations were those that began and ended in the forest state. Those simulations varied on the basis of whether they underwent any transient transitions to quasiperiodicity, and if so, both when the first transition occurred and when the forest state once again became permanent. The difference between the initiation and end of transitions corresponds to one definition of resilience–recovery time, that is, the time taken for a system displaced from equilibrium to return to equilibrium [33]. The static equilibrial systems envisioned in early ecological work is unrealistic, but the definition is transferable to a framework of dynamic equilibria and stable states as presented here.

Resistant systems are of great interest, because although sea urchin populations may increase during outbreaks of SSWD or MHW (Fig. 7A), the community will not transition to barrens. Three conclusions can be drawn from the frequency of resistant responses. First, results fall neatly into three groups. Responses to warming only were overwhelmingly resistant, whereas those perturbed by disease, or disease plus warming, were more likely to be resistant if sea otters were present (Fig. 8). This suggests that warming by the ocean blob during 2013-2015 by itself would not have caused widespread transitions of forests to urchin barrens, but that it did act synergistically with SSWD, perhaps even increasing the probability of the epidemic [43]. Second, both Historical and Modern forests subjected to disease or disease and warming, are more likely to be resistant when sea otters are present (Fig. 8), confirming the keystone role of this species in maintaining giant kelp communities in the forest state. Third, the presence of sea cows confers additional resistance to the system, which we refer to as the “sea cow effect”.

We can now summarize the distinctions between the Historical and Modern communities. Warming alone has minimal effect on forests, Historical or Modern. SSWD alone or with warming, affects almost all the models equally, except for the Historical community with sea otters present. That community would have been up to ten times more likely to undergo transitions to quasiperiodicity, at a rate of 32.23%. However, among resilient systems, any Historical community with or without sea otters would be more resistant to the appearance of barrens at any time during perturbation.

#### 4.4.1 Onset of transitions

When forests did undergo transient transitions to barrens, the questions of interest are when does the first transition occur after the onset of perturbation, and when does the system undergo a final transition back to the forest state, acknowledging that the system may be in the quasiperiodic state during the intervening interval (Fig. 7C).

The onset of the first transition from forest to barrens, in response to SSWD, or warming combined with disease, differed significantly among model-perturbation combinations dependent on two factors: the presence of one or both mammalian species in the system, and the impact of warming. Warming, as explained earlier, indirectly affects several parameters of kelp, algae and echinoderms via its direct impact on metabolic rates. For the Historic system, mean onset time is 7.5 years after the initiation of the perturbation if sea otters are present, which is a considerable delay, and greater than if sea otters are absent–5.9 years. In contrast, if the Modern system is subjected to SSWD only, sea otters have no effect and mean onset time is 6.25 years. This is delayed compared to the Historic system without otters, but more than a year earlier than the Historic system with otters. Sea otters do influence onset time of the Modern system when it is subjected to both warming and SSWD, being 7.35 years, comparable to the Historic system with sea otters present. With the otters absent, mean onset time is 6.45 years, which is delayed compared to the Historic system.

Thus the best of all possible worlds for *M. pyrifera* forest resilience when perturbed by SSWD, or the disease combined with warming, occurs in the presence of both Steller’s sea cow and sea otters. In the present day, without the sea cow, the outcome depends on the perturbation, with the system faring best with sea otters present, and if perturbed by both warming and SSWD. The disease by itself affects the system similarly to the combined effect of warming and disease in the absence of the otter. Although this may appear to be counterintuitive, with a combined perturbation sometimes having a lesser effect than a single one, and also running counter to prior suggestions that warming is a major driver of the transition to urchin barrens [44], a careful consideration of the system offers several synergistic explanations.

Onset time is influenced most by the removal of sunflower sea star predation on the urchins. Urchins thrive under that condition, even in the presence of sea otters. This is not the case when the Modern system is subjected to both warming and SSWD, because urchins are affected negatively by warming and thus do not benefit from predation release to the same extent as when warming is absent. This distinction does not unfold in the Historic systems because Steller’s sea cow moderates the system regardless of the presence or absence of sea otters, or the type of perturbation. Thus, the delay of a transition to urchin barrens when a Modern forest is disturbed is controlled by interactions of the sea urchins with their predators, but the delay of a transition would be enhanced in the presence of the extinct megaherbivore. The presence of sea otters in the Historical system had a negative, delaying effect on the onset of transitions to quasiperiodicity. That effect is not present in the Modern system, highlighting an indirect synergistic interaction between the two mammals that was lost with the extinction of *H. gigas*.

#### 4.4.2 Recovery

Recovery from the last transient barrens state to a permanent forest state does not vary among the models, nor does it differ according to whether the perturbation is SSWD only, or a combination of warming and disease. The mean time to recovery is approximately 9.6 years after the onset of SSWD. The disease itself persists for 10 years in the model, and *P. helianthoides* populations recover rapidly after that. Simulated communities recover earlier than the end of the disease because they are not trapped within the barrens state during the disease, but instead oscillate quasiperiodically between barrens and forest. Thus, if there is a transition to forest within the final year of the disease, the system remains in that state. Kelp forests are beginning to recover in 2022 at various locations in California, nine years after the first appearance of SSWD [20]. Many of these areas are the sites of conservation and restoration efforts that consist of large scale removal of urchins, and in some locations of northern California the re-planting of bull kelp, *Nereocystis luetkana* [17, 45]. The congruent timing of these efforts with the average recovery time predicted by our modeling might be coincidental, but urchin removal does mimic the recovery of sunflower sea star populations [17], and the model supports the timing as being optimal for forest recovery. Filbee-Dexter and Scheibling [7] have suggested that the barrens state may be unstable because of predation, and that is supported by the positive impact of recovering *P. helianthoides* populations in our model. Unfortunately, evidence for actual widespread sea star recovery from SSWD is currently equivocal at best [46]. Nevertheless, our model results, which predict a delayed onset of transitions if Steller’s sea cow is present, also predict that recovery times would have been shorter in the presence of the sea cow.

### 4.5 Restoring functionality and regenerating forests

Predictions of Steller’s sea cow’s impact on kelp forests are varied and depend on the type of perturbation to the system, as well as characteristics of other organisms in the system (e.g., sea urchin ecological efficiency). Could a framework of enhanced kelp forest resilience be implemented in which our model results are operationalized? Functional restoration would consist of artificial mimicry of sea cow browsing by the removal of canopy fronds. Given the complicated nature of the system’s dynamics, the circumstances under which removal is done would depend on the perturbation, as well as the presence/absence of sea otters. Our model predicts a low probability of potential perturbations causing a permanent loss of forest, although it is considerably greater if sea cows were present and the perturbation is a combination of warming and SSWD. One must also account for ongoing changes due to global warming. A more likely outcome under current conditions is the temporary transition to a quasiperiodic state that could persist for at least 10 years. The dynamics of northern Californian kelp forests in the absence of either type of perturbation are not steady state, as shown by multi-decadal records of canopy cover [44]. In fact it could be argued that quasiperiodicity on a timescale longer than predicted by our model might not be uncommon. Both our model and a close examination of canopy cover records [44] suggest that MHW alone may be insufficient to trigger the transitions observed in the past ten years. Instead, a decline in predation pressure and/or a synergy with warming are more likely culprits.

During intervals of prolonged barrens and delays to forest recovery, the probability of demographic or stochastic extirpation of kelp or understory algae increase. The trophic position of these key primary producers would cause such extirpations to cascade through the broader community network, resulting in further biodiversity losses. All these possibilities have been observed in forest-to-barrens transitions. Could our model results be used to increase forest resilience by ensuring greater resistance to transition, and/or reducing recovery times?

The model predicts that the restoration of sea cow functionality in the form of reducing frond canopy cover could be key to both the resistance and resilience of *M. pyrifera* forests when sea otters are absent. Given uncertainty about *H. gigas* consumption rates, however, the intensity of artificial browsing by canopy reduction would have to be established experimentally. Experimental manipulations would also be necessary to assess the roles of other species; for example, the impacts of other urchin predators such as the sheephead *Semicossyphus pulcher* [47]. Finally, such experiments could facilitate the re-direction of kelp productivity back into the forest community. Today, significant amounts of kelp productivity are exported offshore in the form of kelp detritus, where it is believed to comprise a major nutrient supply to the offshore water column [48]. That export would have been less in the presence of the sea cow, and the egestion and excretion of the megaherbivore would have created a significant nutrient feedback directly into the forest community, but that is no longer present. Local harvesting of the kelp canopy could therefore recreate some ecological functionality of the sea cow, increasing forest resilience under certain circumstances while providing an additional nutrient source to the community.

Two important caveats must be considered that are not currently incorporated into our model. First, there could be an onset or recurrence of warming during an outbreak of SSWD, or warming that lasts much longer than the 3 years experienced in the most recent episode and as implemented in the model. Second, urchin populations in our simulations decline as their kelp and understory algal food supplies dwindle, but there is ample evidence that *S. purpuratus* individuals can sometimes persist through years long intervals of starvation at minimal levels of metabolism and body mass–the so-called “zombie urchins” [16]. This persistence could extend kelp and algal recovery relative to our model prediction.

### 4.6 Proposing a new approach

Never before in human history has the need for ecological regeneration been greater than it is today. Yet, the science behind ecological restoration and regeneration remains in its relative infancy. We believe that the Past-Present-Future approach advocated here has potential to be a valuable tool to help close this knowledge gap, enabling researchers to apply a much higher degree of rigor than previously possible, to iterate and track interventions, and thereby enable the field of conservation paleobiology to grow rapidly. Globally, recognition is increasing around the need not only to protect biologically diverse ecosystems, but in many cases to “rewild”, “restore,” and “regenerate” them [49]. Increasingly, conservationists are moving away from a sole focus on endangered species toward collaborative efforts that increase ecosystem abundance, function, and integrity [50]. The current UN Decade on Ecosystem Restoration is a global case in point, with an explicit focus on reviving damaged ecosystems [51]. Recovering ecosystem function is also an explicit aim of the growing array of rewilding projects [52]. At the time of writing, a bill moving through the United States Congress, *Recovering America’s Wildlife Act*, seeks to “provide states, territories, and tribes with $1.39 billion annually to catalyze proactive, on-the-ground, collaborative efforts to restore essential habitat and implement key conservation strategies” [53]; here too emphasis is on the importance of re-establishing biodiversity abundance as an essential step toward ecosystem integrity and resilience. We strongly support this trend to shift the primary focus of conservation from endangered species to ecosystem function. Nevertheless, if these varied and inspiring interventions are to be successful, they must be rooted in rigorous scientific approaches that broaden the time horizon of concern and seek to answer new kinds of questions. What is the current health status (e.g., diversity, resilience, integrity) of the ecosystem in question? What did this ecosystem look like in the past when it was healthier? Given current trends of anthropogenic change, what kinds of interventions, in what order and over what time period, are most likely to generate positive results?

Accurately answering these questions will require a PPF lens that extends the standard time frame of conservation, and utilizes multiple tools and types of data. Conservation paleobiology [1] is the temporal framework for exploring paleoecological snapshots that extend at least as far back as the Pleistocene or early Holocene. With regard to more recent time frames, the traditional ecological knowledge (TEK) of Indigenous cultures has exceptional potential to inject essential information on, for example, shifting trends in species composition and abundance within a given ecosystem [54]. Natural history museum collections offer troves of historical data for understanding past ecosystems and thereby illuminating decisions on conservation interventions. Similarly, publicly-driven community or citizen science efforts and applications (e,g, *iNaturalist*, *eBird*), are continuously generating rich data about present day (and recent past) biodiversity occurrences [55]. Moving forward, community science efforts will be increasingly needed to monitor the impacts of conservation actions, generating data that can be used to improve ecological models. Could similar efforts be designed for the paleontological record?

Mathematical modeling, which is at the core of this study, has exceptional potential to interweave these diverse data streams [56] and, in doing so, provide answers to the above questions. Models offer the advantage over strictly qualitative or data-analytical approaches of being capable of rigorous exploration of future or counterfactual ecosystem states. The methodology advocated here can be summarized as follows. First, establish basic ecological networks (matter and energy flows) of a present-day ecosystem, including both environmental conditions and biological components. Second, integrate historical and/or paleontological data into the model to explore putative past states of that ecosystem, distilling key elements, such as megaherbivore impacts and predator-prey relationships that no longer exist. Third, integrate the first and second steps to create mathematical models of future ecosystem states, exploring different state iterations under varying environmental conditions and key taxon compositions. Finally, apply the latter to generate predictions about the likely impacts of specific interventions, and use these predictions to guide experimentation, policies, and action.

## 5 Summary

As anthropogenically-driven environmental changes multiply, intensify and become more permanent features of the natural world, we will need an increasingly diverse and powerful set of tools to address the conservation and regeneration of natural systems. Similarly, as species decline, and extirpations and extinctions increase, historical and paleontological data and analyses must be brought to bear in efforts to understand the altered and no-analog states of modern ecosystems. In this quest, we believe that the PPF mathematical modeling approach advocated in this paper has considerable potential. Here we combined paleontological, historical and ecological data into a detailed mathematical model of how an extinct marine megaherbivore, Steller’s sea cow, might have affected the dynamics and resilience of an important coastal marine ecosystem. Those effects were generally non-trivial and often counterintuitive. On one hand, the sea cow may have increased the probability that giant kelp forests would switch to a permanent alternative state exhibiting periodic urchin barrens when subjected to marine heat waves and sea star wasting disease. Yet, on the other hand, forests that did not transition to an alternative state would have been significantly more resilient–either resisting even transient transitions or recovering from them more quickly than today’s forests. Moreover, when both sea cows and sea otters were present in historical forests, they would have interacted indirectly to delay the onset of transitions after the initiation of the combined perturbations, a function that the sea otters today apparently do not fulfil.

The rapid degradation and, in some cases, collapse of modern ecosystems has spurred actions that go beyond traditional species conservation, instead focusing on active stewardship and regeneration of those systems. For example, wildfires in western North America that have been growing in frequency, intensity and multi-seasonality, are being increasingly addressed with management techniques that intervene directly in forest composition and dynamics, such as tree thinning and prescribed burns [57]. Those techniques in turn are informed by ecological models [58], field experimentation [59], as well as deep Indigenous histories of active landscape management [60, 61]. Similar interventions are largely absent from marine systems, with efforts focused instead on species conservation and restoration via managed harvesting and marine protected areas. Active interventions that have been suggested–such as the large-scale addition of supplemental iron to open ocean surface waters to spur increased phytoplanktonic productivity to absorb more atmospheric carbon dioxide– have rightfully been treated as highly speculative and potentially dangerous given our limited understanding of the complex pathways and feedbacks involved [62]. Here we propose an intervention based on a detailed mathematical model, one that seeks to restore a natural ecosystem function that was a part of the giant kelp ecosystem in the relatively recent past. Any such intervention, however, would have to be preceded first by extensive *in situ* experiments to both enhance model predictions with better model parameterization, and to identify important components that may currently be absent from the model. Nevertheless, in our view the dire state of giant kelp ecosystems justify consideration of this approach.

There is a growing appreciation of the roles of extant and extinct megafauna as ecosystem engineers [63]. Direct impacts of large vertebrates include physical habitat restructuring, consumption of large quantities of biomass at or near the base of food chains, facilitation of processes vital to the survival of other species, and flux rates and transport of nutrients. Examples include tree destruction by foraging and moving elephants [64], planktivory by baleen whales [65], germination of seeds by large mammals and birds such as gomphotheres and dodos [66, 67], and both whale falls [68] and benthic feeding by grey whales [69]. To this may be added the browsing of giant kelp canopies by Steller’s sea cow. Important functions of megafauna may extend well beyond such direct impacts, however, and must include the indirect interactions and impacts of these powerful consumers that affect community resilience and alternative state dynamics. Efforts to conserve and regenerate natural systems can, in our opinion, benefit from consideration of novel, mathematically rigorous interventions like the one proposed here.

Humans are fully interdependent with Earth’s ecosystems, and thus our fates are deeply intertwined. Together, the initiatives listed above (and many others) could be transformative, ultimately with potential to help redefine the relationship between humans and nonhuman nature.

## Author Contributions

RB, PR and SS conceived of the study. PR developed the mathematical model and wrote simulation code. RB and PR conducted statistical analyses. All authors contributed to the final manuscript and approved the final submission.

## Acknowledgments

The authors acknowledge that the California Academy of Sciences occupies land that is the ancestral homeland of the Ramaytush, the vast majority of whom were forcibly removed from their homelands. We acknowledge this painful history and honor their legacy and continuing contributions to better understanding the natural history of the United States.

## Supplementary material

### 1 Extinction hypotheses

Several hypotheses have been put forward regarding the extinction of *H. damalis*. It is certain that pre-industrial hunting is responsible for the death of the last individuals, but the single known population was geographically restricted, and may have numbered as few as 2,000 individuals [1]. Recent reconstruction of a *H. damalis* genome points to a population bottleneck suffered roughly 400,000 years ago. Coupling the subsequent reduction of genetic diversity with presumed population fragmentation during intervals of glacially-lowered sea level, Sharko et al. [1] suggested that the species was doomed to extinction prior to historical exploitation, a type of “dead clade walking” [2]. There are arguments that the lowering of sea level during intervals of glacial expansion would have reduced the shallow coastal habitat required by giant kelp, and the Comander Islands population was therefore a relict one. The Comander Islands, however, are not the likeliest location for a glacial refuge in the North Pacific, because of the limited benthic shelf area that would be available for giant kelp forests. Significantly more habitat space would have been available in the Bering Sea and off southeast Alaska.

It remains possible though that the Comander Islands population was itself the remnant of a recent expansion following the last glacial termination approximately 10,000 years ago. Graham et al. [3] have argued for a threefold expansion of *M. pyrifera* forests in southern California between the last glacial maximum and the mid-Holocene, followed by a rapid decline. This type of response probably also occurred further to the north [3], although the timing may have differed. In that case, the historical population of the Comander Islands may indeed have been a geographic relict, but the ability of the species to undergo range expansions and contractions, and to persist through multiple glacial-interglacial cycles, would refute the hypothesis that this was a clade pre-destined for extinction. Our argument is supported by the discovery of *H. gigas* bones from St. Lawrence Island to the north in the Bering Sea, dated to 800-920 CE.

Hunting by regional Indigenous peoples has also been proposed as a cause of the species’ limited range by historical times, although there is no archaeological evidence to support this. Crerar et al. [4] suggested that Inuit, expanding into the Bering Sea during the Medieval Warm Period (950-1250 CE), may have hunted the species, basing this on the coincidental occurrence of Inuit migration with the St. Lawrence Island specimen, but there are no records of hunting, and no remains of *H. gigas* have to-date been associated with the activities of Indigenous peoples.

Finally, it has been proposed that the small population size and geographic restriction documented by Steller were secondary consequences of a widespread transition of giant kelp forests to urchin barrens caused by Indigenous and Russian over-exploitation of sea otters [5, 6, 7]. There is no evidence that Indigenous peoples over-exploited sea otters to the extent that kelp forests underwent state transitions. Whether they could have driven population declines or extirpation of *H. gigas* is an open question. What is entirely plausible is that *H. gigas*, like many other Pleistocene-Holocene mammalian megafaunal species may have been vulnerable to the interval’s significant climatic fluctuations, and thus to the combined climate change and prehistoric hunting proposed for terrestrial scenarios in the Americas and Australia [8, 9, 10]. Population modeling of *H. gigas* by Turvey and Risley [11] supports the contention that small bands of pre-industrial hunters could have been a sole cause of extirpation, but that the documented intensity of 18^th^ century hunting suggests that the Comander Island population was actually larger than indicated in written accounts. Therefore the only certainty is that the last known individuals of Steller’s sea cow were killed and consumed by Russia-based commercial venturers.

### 2 Supplementary Methods

#### 2.1 Physical parameters

Three physical parameters are present in the models: seasonal day length, which controls kelp and understory algal productivity; wave energy, which creates the probability that adult kelp will be dislodged and removed from the population; and sea surface temperature, which seasonally affects the metabolic rates and therefore ecological parameters of kelp, understory algae, and invertebrates.

#### 2.1.1 Temperature

A core tenet of the Metabolic Theory of Ecology (MTE) is that environmental temperature affects the basal cellular metabolic rates of many organisms [12], and as a consequence, the efficiency of energy assimilation and intrinsic rates of increase (“*r*”) of species populations. For example, a typical relationship exists between maximum *r*, designated here as *r*_max_, at an optimum temperature *T*_opt_ (other physical factors being held constant). *r* declines slowly below *T*_opt_, often remaining positive down to 0^°^C for organisms in cool temperate and polar waters [13]. In contrast, *r* may decline sharply above *T*_opt_, approaching zero between 20-25^°^C for those same organisms. Empirical determinations of the sea surface temperature (SST)-*r* relationship are few, but resemble inverted skewed parabolas and are generally fit with a Sharpe-Schoofield model [14]. Here we model the relationships using a Linex loss function [15], which bears a similar geometric form. For example, the dependence of *r* on temperature is described as

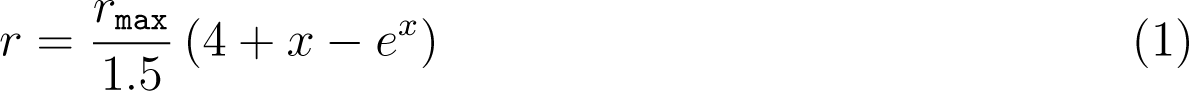

where

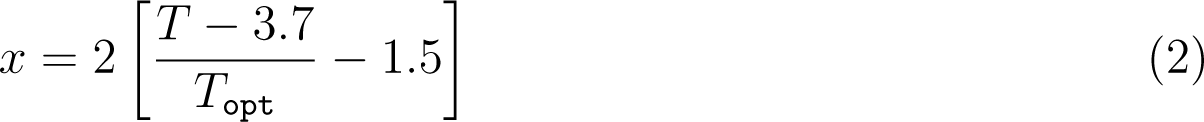

The choice of this function is *ad hoc*, but it captures the essential morphology of the relationship (Fig. 1A), including positivity at 0^°^C, a maximum *r* at *T*_opt_, and an intercept of the SST axis at *≈* 25^°^C. *T*_opt_ itself is determined either from empirical studies of the model system’s species, or from the maximum seasonal temperature of the mid-point of the geographic range of a species for which the value is unmeasured. The occurrences and densities of both *S. purpuratus* and *P. helianthoides* have been shown to be temperature-dependent, with *S. purpuratus* being increasingly common at temperatures exceeding 14^°^C and *P. helianthoides* less common [16]. Model optimal temperatures are listed in Table 1 for *M. pyrifera*, *C. corymbiferus*, *S. purpuratus* and *P. helianthoides*.

Sea surface temperature itself is calculated as

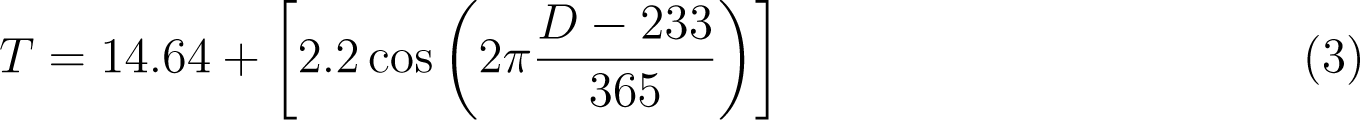

where *D* is the numerical day of the year, and 14.64*°*C is the mean annual SST of Monterey Bay, California (Monterey Bay National Marine Sanctuary) (Fig. 1B). This function has a low frequency, multi-decadal scale variation that captures decadal variation in the North Pacific, though this is not relevant for the sub-decadal perturbations considered in this study.

#### 2.1.2 Day length

Day length Δ*d* is the number of hours per day for which light intensity is above the minimum threshold required for photosynthesis (*≈*1%). It varies latitudinally and seasonally. Annual daylength is calculated for Monterey Bay based on a maximum of 14.67 hours of daylight and a minimum of 9.67 hours, as

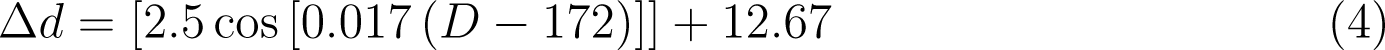

where *D* is the numerical day of the year (Fig. 1C).

#### 2.1.3 Hydrodynamic forces

On the central California coast today, the major source of kelp mortality in the forested state is dislodgement by hydrodynamic forces. Kelp can withstand substantial drag because of their flexibility, and the attenuation of wave energy by the density of the forest itself. Nevertheless, high energy waves and storm surges are capable of dislodging giant kelp. Most of this mortality occurs in late winter, when up to 90% of adult individuals in a single forest can be removed [17]. This process is likely very important to the community’s dynamics as it both provides a mechanism for persistence of understory algae, as well as creates a decline of the drift detritus supply.

We modeled storm occurrence and associated wave height based on sub-daily wave height measures in Monterey Bay for the years 2009-2020 (National Oceanographic and Atmospheric Administration, Monterey Bay Aquarium historical sensor data, Buoy Station - sea surface wave significant height). The model simplifies the fact that any single event of extreme wave height is likely to have an autocorrelation length of several days, where the occurrence of extreme waves is distributed over several consecutive days. This means that a cumulative total of *n* kelp plants are removed during that entire interval, not *n* plants per day. Combining multiple years of data can exacerbate this effect by increasing the possible duration of events because they occur at approximately the same times of year, but not exactly on the same days. Storm surge events were therefore sampled from a modeled smooth cosine function that captures, at maximum amplification, the wave height threshold above which adult kelp become susceptible to removal. To this function is added an uncorrelated daily noise, tuned low enough to allow persistence of the biotic system (Fig. 1D).

Wave height was therefore modeled as a wave that has its maximum amplitude at 2.5 m, to which is added a noise sufficient to capture stochastically extreme wave heights sufficient to dislodge kelp.

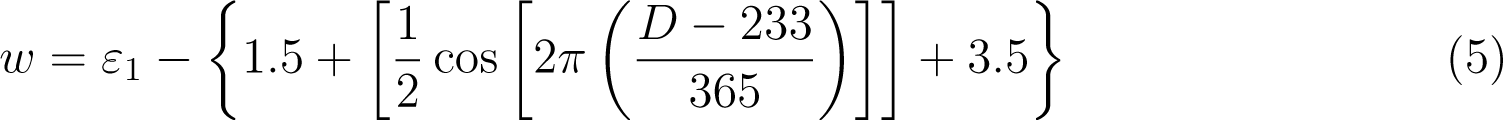

where *w* is wave height (meters) and *η*_1_ is a random noise ranging [*−*0.1, 0.1]. Furthermore, *δ_M_*, the surge-driven mortality rate of adult kelp, is given as

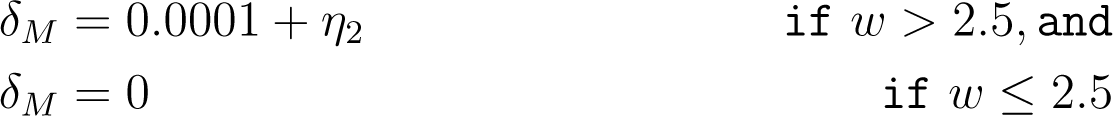

where *η*_2_ is a random noise ranging (0, 0.000075].

### 2.2 Probability of sea otter presence

Sea otters (*Enhydris lutris*) are present in a community only when sea urchins (*Strongylocentrotus purpuratus*) are of sufficient nutritive value. That value declines as a function of declining kelp and other edible algal densities, and is typically measured as gonadal index, which is the fraction of body weight accounted for by gonadal tissues. Smith et al. [18] related the frequency of sea otter site selection for predation to gonadal index. From their data (their Fig. 4) we estimated the following relationship between the probability of otter predation and gonadal index

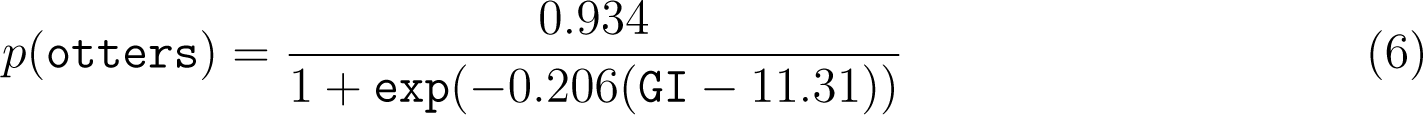

where GI is gonadal index. GI in turn is derived by inverting the relationship between it and the fraction of the urchin population that is exposed (given by text Equation 17),

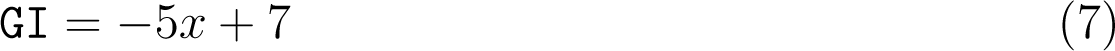

### 2.3 Model parameterization

Model parameters were determined using a combination of both empirical and theoretical relationships among organismal traits and external variables, and a criterion of system feasibility; that is, all species persist and co-exist in an unperturbed community and none become extinct during the burn-in interval. A complete list is given in Table 1. Most parameter values were invariable among models and simulations, but several were drawn stochastically from distributions to reflect real-world uncertainty and variability of their values. The latter include: wave height; the presence of sea otters as urchin predators; rates of intrinsic increase of the algae *M. pyrifera* and *C. corymbiferus*; consumer mortality rates; and ecological efficiencies of the consumers *S. purpuratus*, *P. helianthoides* and *H. gigas*.

### 2.3.1 Rates of intrinsic increase

Rates of intrinsic increase of *M. pyrifera* and *C. corymbiferus* were set at 0.1*±λ*, where *λ* is drawn randomly from a normal distribution of mean zero and standard deviation 0.025 (*N* (0, 0.025)).

### 2.3.2 Algal recruitment rates

Rates of recruitment of *M. pyrifera* and *C. corymbiferus* were assumed to be 0.001d*^−^*^1^ and 0.0001d*^−^*^1^ respectively. These rates are derived from those presented by Detmer et al. [19], but modified to maintain feasibility of the populations under the conditions of both spatial competition and predation in the current model.

### 2.3.3 Kelp frond senescence

Kelp fronds eventually decline in photosynthetic activity and detach from the main stipe, becoming part of the pool of drift detritus. They do this in the model at a rate of 0.0125d*^−^*^1^ [19].

### 2.3.4 Rates of mortality

Mortality rates of the consumers (otters excepted as their demographics are not controlled by model interactions) were estimated based on empirical scaling relationships between measured mortality rates and body masses for a wide range of organisms [20]. The relationship was further scaled to yield feasibility of more than 90% of unperturbed models, as determined by 100 simulations per model type (Historical and Modern, with and without otters). The relationship is

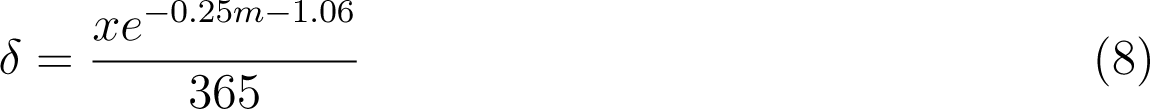

where *δ* is a daily mortality rate, *m* is body mass (grams), and *x* =300 for *S. purpuratus* and 30 for *P. helianthoides* and *H. gigas*.

### 2.3.5 Ecological efficiency

Ecological efficiency, *ε*, is the rate or efficiency with which consumed prey are converted into new predators. Estimates vary widely in theoretical models, but are often assumed to be 0.1. This value is justified as the thermodynamic “rule of thumb” of an *≈*10% efficiency of energy assimilated from consumed organic material. That value is used here for sea urchin, sunflower sea star and sea cow efficiencies, but to each value of 0.1 per simulation is added a small stochastic variation drawn from a normal distribution, *N* (0.0,0.003).

### 2.3.6 Otter predation

The probability that otters are present and preying on *S. purpuratus* was derived from an empirical study of otter predation and urchin nutritive condition as measured using gonadal index [18]. Because urchin condition is a function of kelp density, we related otter activity directly to kelp density (text Eq. 17). We rendered the relationship probabilistic by having otters present if *E < p*, where *p* is generated randomly from a uniform distribution.

## 3 Supplementary Tables and Figures

### 3.1 Tables

**Table 1:** Variable and parameter definitions and values.

| Parameter | Definition | Value |
| --- | --- | --- |
| $r_{\max}$ | Maximum rate of intrinsic increase of *M. pyrifera* and *C. corymbiferus* | 0.1 day<sup>-1</sup> |
| $T_{\text{opt}}$ | Temperature at which $r$, $\alpha$ or $\varepsilon$ is maximum for a species | |
|  | *M. pyrifera* | 15°C |
|  | *C. corymbiferus* | 13°C |
|  | *S. purpuratus* | 16°C |
|  | *P. helianthoides* | 13°C |
| $D$ | Numerical day of the year | 1-365 |
| $\eta_1$ | Random noise added to deterministic wave height, $w$ | [-0.1, 0.1] |
| $\eta_2$ | Random noise added to wave-driven *M. pyrifera* mortality, $\delta_M$ | (0, 0.000075] |
| $\sigma_M, \sigma_C$ | *M. pyrifera* and *C. corymbiferus* recruitment rates | 0.001 day<sup>-1</sup> |
| $\alpha_i$ | Predator-prey interaction coefficients | |
| $\alpha_1$ | *S. purpuratus* - *M. pyrifera*, *C. corymbiferus* | 0.1 |
| $\alpha_2$ | *P. helianthoides* - *S. purpuratus* | 0.05 |
| $\alpha_3$ | *E. lutris* - *S. purpuratus* | 0.00005 |
| $\alpha_4$ | *H. gigas* - *M. pyrifera* fronds | 0.00002 |
| $\delta_F$ | *M. pyrifera* frond senescence rate | 0.0125 day<sup>-1</sup> |
| $\delta_i$ | Mortality rates | |
| $\delta_1$ | *S. purpuratus* | 0.0900 day<sup>-1</sup> |
| $\delta_2$ | *P. helianthoides* | 0.0034 day<sup>-1</sup> |
| $\delta_3$ | *H. gigas* | 0.00005 day<sup>-1</sup> |

**Table 2:** Correlation coefficients and probabilities of significance between population parameters and system state.

| Parameter | $r^2$ | p(significance level) |
| --- | --- | --- |
| kelp $r$ | 0.0584 | 0.5639 |
| understory algal $r$ | 0.1372 | 0.1735 |
| sea urchin $\varepsilon$ | 0.7180 | <0.0001 |
| sunflower sea star $\varepsilon$ | 0.0792 | 0.4336 |

**Table 3:** Frequencies of transitions of forested state in response to perturbation. MHW - marine heat ave; SSWD - sea star wasting disease. Frequencies report the number of simulations out of 100 that transitioned to quasiperiodicity, were resistant to transition, or were resilient, instead transitioning temporarily to quasiperiodicity.

| Model | Perturbation | Quasiperiodic | Resistant | Resilient |
| --- | --- | --- | --- | --- |
| Historical | MHW | 2 | 82 | 1 |
|  | SSWD | 5 | 66 | 13 |
|  | MHW+SSWD | 12 | 51 | 20 |
| Historical,<br>no otters | MHW | 0 | 69 | 2 |
|  | SSWD | 2 | 24 | 51 |
|  | MHW+SSWD | 0 | 20 | 49 |
| Modern | MHW | 2 | 68 | 1 |
|  | SSWD | 3 | 45 | 33 |
|  | MHW+SSWD | 1 | 43 | 32 |
| Modern,<br>no otters | MHW | 0 | 67 | 2 |
|  | SSWD | 2 | 16 | 48 |
|  | MHW+SSWD | 0 | 25 | 51 |

**Table 4:**
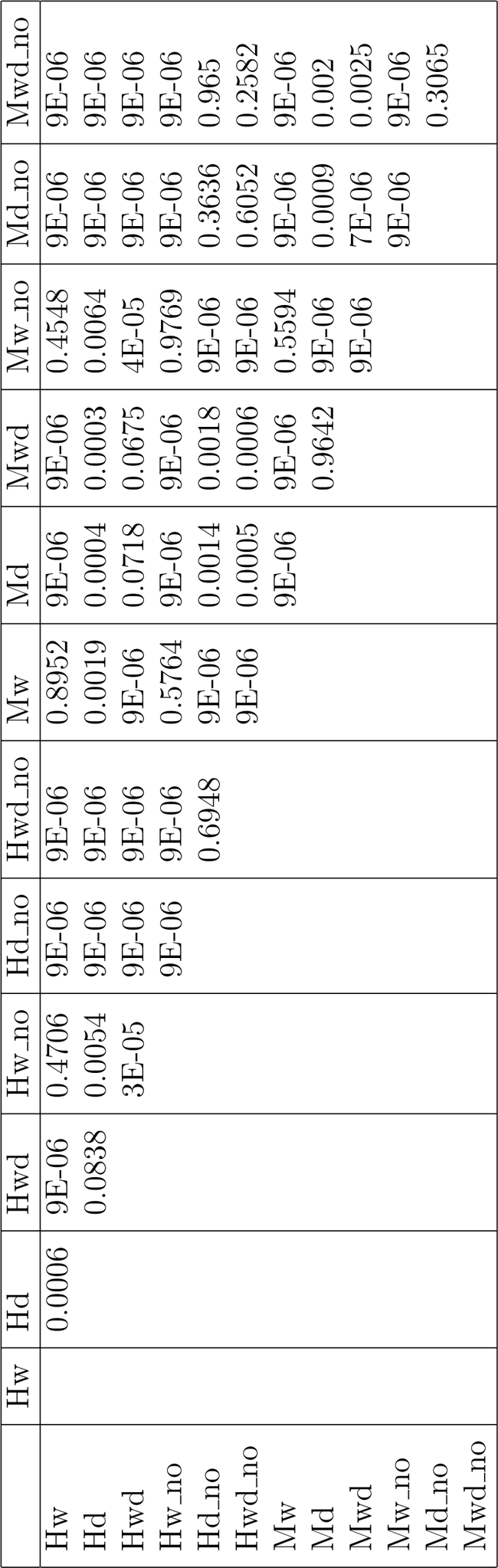
Chi-square probabilities of differences of frequencies of resistant simulations between pairwise comparisons of model-perturbation combinations. See Table S1 for explanation of which models of each pair had the higher (lower) frequency.

**Table 5:** Principal components, principal components analysis of population densities of resilient simulations one year prior to perturbation.

| PC | Eigenvalue | % variance explained |
| --- | --- | --- |
| 1 | 1.796 | 44.90 |
| 2 | 1.129 | 28.23 |
| 3 | 0.600 | 15.01 |
| 4 | 0.474 | 11.86 |

**Table 6:** Principal component loadings of populations on first two principal components of analysis of population densities of resilient simulations one year prior to perturbation.

| Population | PC1 | PC2 |
| --- | --- | --- |
| kelp | 0.5513 | -0.3808 |
| understory algae | -0.4123 | 0.6469 |
| sea urchins | 0.4907 | 0.4944 |
| sunflower sea stars | 0.5341 | 0.4383 |

### 3.2 Figures

**Figure 1:**
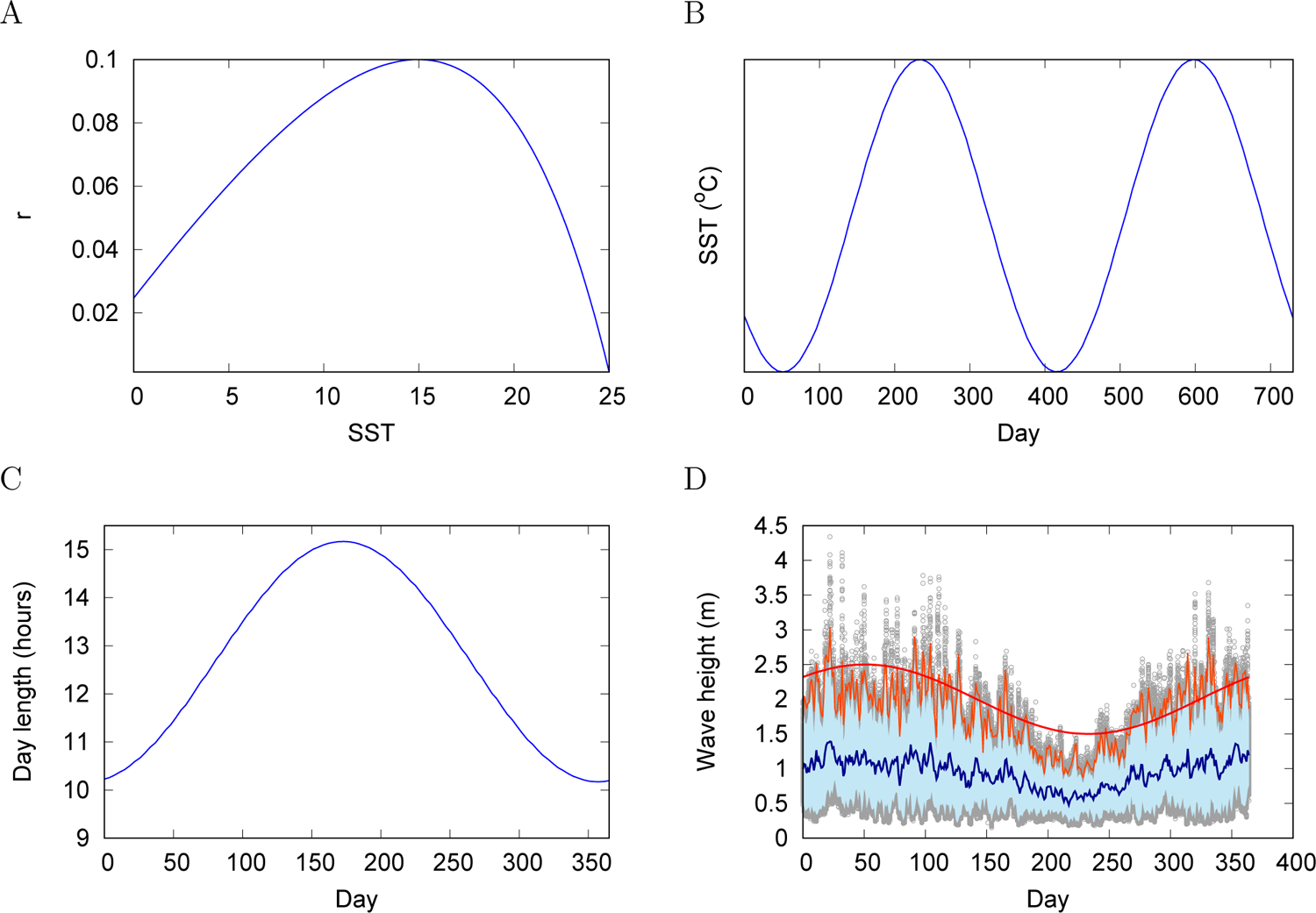
(**A**) Relationship between sea surface temperature (SST) and *r* for the kelp *Macrocystis pyrifera*. *T*_opt_ is 15^°^C. (**B**) Annual SST variation, Monterey Bay. (**C**) Day length, *D*, throughout the year, Monterey Bay. Here *D* refers to light levels sufficient for photosynthesis to occur within the upper 1.0 m of the water column. (**D**) Annual wave height variation, Monterey Bay, 2009-2020. Grey circles - raw data. Dark blue line - mean. Light blue solid background - 5-95% empirical variation. Light red line - approximate daily maximum during total interval, 2.33*σ* above the mean. Solid red line - cosine function estimating daily maxima.

**Figure 2:**
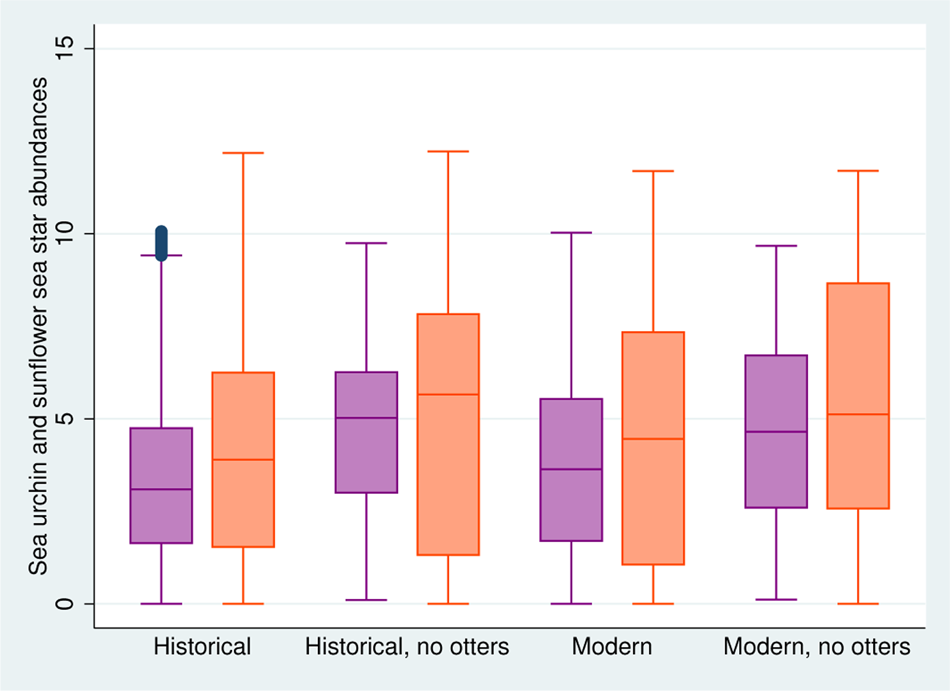
Sea urchin and sunflower sea star relative abundances in unperturbed forests. Purple - sea urchins; orange - sunflower sea stars. There are no differences among model types, but abundances are higher when sea otters are absent.

**Figure 3:**
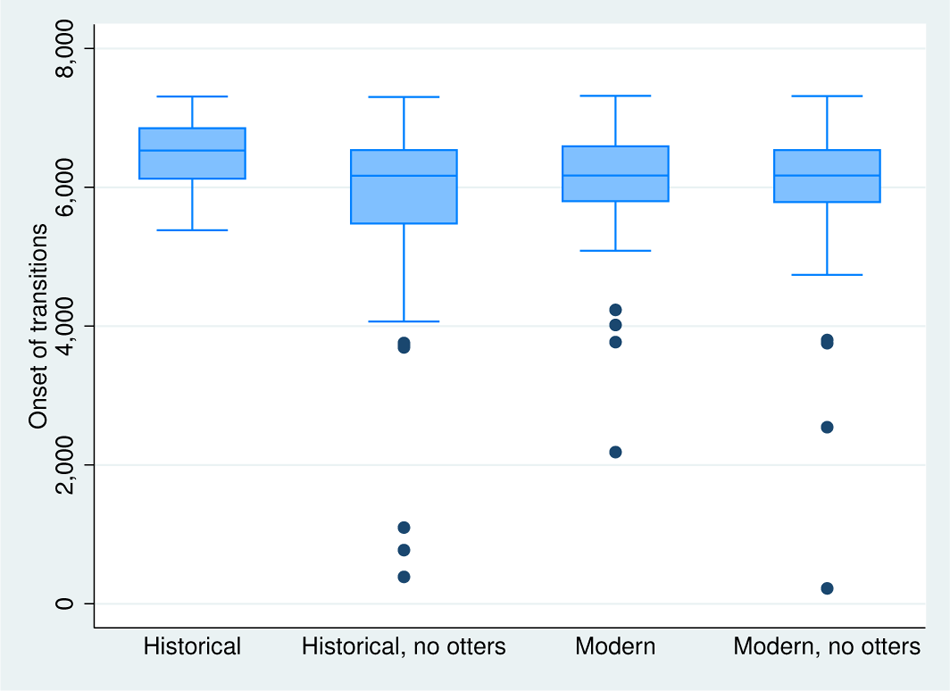
Distributions of onset times of transitions after the initiation of perturbation (day 0). SSWD and MHW+SSWD perturbations are pooled within models.

